# MicroRNA156 and its targeted *SPL* genes interact with the photoperiod, vernalization, and gibberellin pathways to regulate wheat heading time

**DOI:** 10.1101/2025.11.05.686864

**Authors:** Qiujie Liu, Lili Zhang, Zhicheng Zhou, Chaozhong Zhang, Chengxia Li, Juan M. Debernardi, Jorge Dubcovsky

## Abstract

Heading time has a large impact on adaptation to different environments and crop productivity. In this study, we characterized the effect of the endogenous age pathway on heading time and its interactions with the photoperiod and vernalization pathways in the leaves of tetraploid wheat (*Triticum turgidum* ssp. *durum*). Plants with reduced levels of microRNA156 or increased expression of its downstream targets, the *SQUAMOSA PROMOTER BINDING PROTEIN-LIKE* genes *SPL3, SPL4,* and *SPL13* exhibited accelerated heading time, with stronger effects under suboptimal inductive conditions. Earlier heading was associated with the upregulation of miR172 and flowering-promoting genes *VRN1*, *FUL2*, and *FT1* and the downregulation of flowering-repressing genes *AP2L1* and *VRN2*. Additionally, we uncovered complex interactions among SPL, SQUAMOSA (VRN1 and FUL2) and DELLA proteins that modulate wheat heading time. We showed that DELLA proteins, which are negative regulators in the gibberellic acid pathway, can interact with SPL proteins reducing their ability to induce flowering. We also discovered previously unknown interactions between SQUAMOSA and DELLA proteins in wheat that compete with the DELLA-SPL interactions, likely reducing DELLA’s ability to repress SPL3 and SPL4 activity. Since SPL3 and SPL4 directly promote *VRN1* and *FUL2* transcription, these interactions generate a positive regulatory feedback loop that accelerates wheat heading time. Finally, we developed dominant miR156-resistant alleles *rSPL3, rSPL4,* and *rSPL13* that accelerate wheat heading time under both optimal and suboptimal inductive conditions. These publicly available genetic resources can be used to fine tune heading time and improve wheat adaptation to changing environments.

**Significance statement:** Wheat heading time is critical for adaptation to diverse environments. We generated dominant mutations for the *SPL3*, *SPL4* and *SPL13* genes that accelerate heading time. Different combinations of these mutations can be used to modulate heading time and improve wheat adaptation to changing environments.

## Introduction

Wheat (*Triticum aestivum*) provides more than one fifth of the calories and protein consumed by the human population (FAOSTAT, 2023) and is important for global food security. Continuous increases in wheat grain yield are required to feed a growing human population in a rapidly changing environment. An important determinant of wheat adaptation to these changes is the adjustment of the reproductive phase to the time of the year with optimal conditions for reproductive success. The precise timing of the transition from the vegetative to the reproductive phases and of spike emergence (heading time) is critical to maximizing wheat grain yield.

Natural variation in heading time is mainly associated with genes regulating vernalization (long exposures to cold temperatures) and photoperiod (length of the day-night cycle) requirements (Yan *et al*., 2004a, Yan *et al*., 2004b, Fu *et al*., 2005, Yan *et al*., 2006, Beales *et al*., 2007, Distelfeld *et al*., 2009, Wilhelm *et al*., 2009). Wheat cultivars are classified as winter or spring types based on their vernalization requirements, and as photoperiod-sensitive (PS) or photoperiod-insensitive (PI) based on the difference in heading time between long (LD) and short days (SD).

Winter wheat is sown in the fall, when flowering is prevented by high expression levels of the *VERNALIZATION2* (*VRN2)* gene, a LD repressor of the florigen-encoding gene *FLOWERING LOCUS T1* (*FT1*, also known in wheat as *VRN3*) (Yan *et al*., 2004b, Yan *et al*., 2006). The long exposure to cold temperatures during winter results in the upregulation of *VERNALIZATION1* (*VRN1*) in the shoot apical meristem (SAM), which promotes the transition to the reproductive phase (Yan *et al*., 2003, Oliver *et al*., 2009). *VRN1* is also upregulated in the leaves, where it prevents the upregulation of *VRN2* in spring by directly binding to its promoter (Chen and Dubcovsky, 2012, Deng *et al*., 2015). As day length increases during spring, *VRN2* is repressed by *VRN1*, and the *PHOTOPERIOD1* (*PPD1*) gene induces the expression of *FT1* in the leaves. The FT1 protein is then transported to the SAM via the phloem, where it further upregulates *VRN1* expression and promotes GA biosynthesis, thereby accelerating spike development and inducing stem elongation (Pearce *et al*., 2013). A positive feedback loop between *VRN1, VRN2*, and *FT1* in the leaves ensures an irreversible commitment to flowering after vernalization (Loukoianov *et al*., 2005, Distelfeld *et al*., 2009).

Wheat heading time is also regulated by the endogenous age pathway (Debernardi *et al*., 2022). Micro RNA156 (miR156), which is expressed at high levels in juvenile tissues and declines with plant age, plays a central and conserved role in regulating the transition between the juvenile and adult stages during the vegetative growth phase (Lawson and Poethig, 1995, Wu and Poethig, 2006, Poethig, 2009, Wu *et al*., 2009). Constitutive expression of miR156 leads to a prolonged juvenile phase and late flowering, while reducing miR156 activity by expressing artificial target mimics (MIM156) reduces juvenile traits and accelerates flowering (Schwab *et al*., 2005, Xie *et al*., 2006, Chuck *et al*., 2007, Wu *et al*., 2009, Wang *et al*., 2015b, Debernardi *et al*., 2022)

In Arabidopsis, miR156 affects plant development by controlling the expression of *SQUAMOSA PROMOTER BINDING PROTEIN* (*SBP*)-*LIKE* (*SPL*) genes (Wu and Poethig, 2006). Members of this gene family encode plant-specific transcription factors (TFs) that bind to regulatory regions of MADS-box genes within the *SQUAMOSA* clade (Huijser *et al*., 1992, Klein *et al*., 1996) and promote flowering in Arabidopsis (Yu and Wang, 2020). *SPL* genes also promote the transition to the reproductive phase by inducing the expression of miR172 (Wang *et al*., 2009, Wu *et al*., 2009, Wang, 2014, Wang *et al*., 2015a, Hyun *et al*., 2016). The age-dependent induction of miR172 results in the downregulation of its targeted *APETALA2-LIKE* (*AP2L*) genes, which function as repressors of the flowering transition. In wheat and other plant species, ectopic expression of miR172 or knock-out mutants of *AP2L* genes accelerate flowering, whereas transgenic expression of a target mimicry against miR172 (MIM172) or miR172-resistant versions of *AP2L* genes (*rAP2L*) delay flowering (Aukerman and Sakai, 2003, Lee *et al*., 2014, Debernardi *et al*., 2022). The expression of miR172 is also regulated by *GIGANTEA* in both Arabidopsis (Jung *et al*., 2007) and wheat (Li *et al*., 2024), linking the endogenous and photoperiod pathways.

The activity of some SPL transcription factors is also regulated through protein-protein interactions with DELLAs (Yu *et al*., 2012), which are negative regulators in the gibberellin (GA) signaling pathway. In the absence of GA, DELLA proteins repress growth through physical interaction with their target proteins (Sun, 2010). GA promotes the degradation of DELLA by binding to a nuclear receptor, GIBBERELLIN INSENSITIVE DWARF1 (GID1) (Griffiths *et al*., 2006, Harberd *et al*., 2009, Sun, 2010). The roles of DELLA proteins and GA in the regulation of the SPL activity and the duration of the juvenile phase contribute to their effects on flowering time in Arabidopsis (Galvao *et al*., 2012, Yu *et al*., 2012, Hyun *et al*., 2016). In the absence of GA, DELLA can directly interact with SPL9 and interfere with its transcriptional activity (Yu *et al*., 2012). Increasing levels of GA with plant age promote DELLA degradation, thereby releasing SPL proteins that promote flowering. GA-deficient mutants or mutants impaired in GA signaling show late flowering, whereas exogenous GA application in wheat accelerates heading time (Griffiths *et al*., 2006, Pearce *et al*., 2013, Wang, 2014).

We have recently shown that in spring wheat the miR156-miR172-AP2L pathway regulates *FT1* expression in the leaves, whereas in winter wheat the regulation of *AP2L1* expression is decoupled from the age-dependent downregulation of miR156. In winter varieties, the induction of *VRN1* by vernalization is required to repress *AP2L1* in the leaves and promote flowering (Debernardi *et al*., 2022). However, the roles of wheat *SPL* genes in the endogenous age pathway and their interactions with the distinctive photoperiod and vernalization pathways of temperate grasses remain underexplored.

In this work, we identified three wheat *SPL* genes regulated by miR156 in the leaves and characterized their effects on heading time under optimal and sub-optimal photoperiodic and vernalization conditions. We also explored the interactions among the endogenous age pathway and the photoperiod, vernalization, and GA pathways and discovered interactions between DELLA and SQUAMOSA proteins that have not been reported before. Finally, we developed dominant *SPL* alleles and discussed how they can be manipulated to fine-tune wheat heading time and facilitate the development of wheat varieties better adapted to changing environments.

## Materials and Methods

The phylogenetic analysis of wheat (*Triticum aestivum*), rice (*Oryza sativa*) and Arabidopsis SPL proteins based on their SBP domains is described in Method S1. Total RNA and small RNA extraction and purification, as well as qRT-PCR quantification protocols, are described in Method S2. Plant materials and mutants used in this study, and their growing conditions are described in Supplemental Method S3. The methods used to develop the *SPL13* CRISPR mutants and the transgenic lines overexpressing this gene are described in Method S4. The methods used to characterize DNA-protein interactions included Method S5 for yeast-one-hybrid assays (Y1H) and Method S6 for electrophoretic mobility shift assays. Methods for protein-protein interaction, including yeast-two-hybrid assays (Y2H), bimolecular fluorescence complementation (BiFC), coimmunoprecipitation (Co-IP), and yeast-three-hybrid assays (Y3H) are described in Methods S7, S8, S9, and S10, respectively. Statistical methods are summarized in Method S11.

## Results

### Identification of miR156-targeted SPL genes in tetraploid wheat

We identified nineteen *SPL* genes in each of the three genomes of hexaploid wheat (*Triticum aestivum*, genomes AABBDD) cultivar Chinese Spring (International Wheat Genome Sequencing Consortium, 2018) (RefSeq v1.1), nine of which have complementary regions to miR156 (*SPL2*, *SPL3*, *SPL4*, *SPL7*, *SPL13*, *SPL14*, *SPL16*, *SPL17*, and *SPL18*, Figure S1a).

Using an alignment of the conserved SBP domains from wheat, rice, and Arabidopsis SPL proteins (Figure S2), we generated a phylogenetic tree (Figure S3, Method S1), which is similar to previous studies (Cao *et al*., 2019, Cao *et al*., 2021, Chen *et al*., 2023, Gupta *et al*., 2023). *SPL* genes were named (Data S1) following a published phylogenetic-based wheat nomenclature (Gupta *et al*., 2023).

Using previously published RNA-seq data from multiple mature tissues (Figure S1b) and single-molecule fluorescence in situ hybridization (smFISH) in the early transition of the SAM to an inflorescence meristem (IM, Figure S1c) (Xu *et al*., 2025), we found that among the nine *SPL* genes with miR156 binding sites (Figure S1a), four were expressed in mature leaves and developing spikes (*SPL2*, *SPL3, SPL4,* and *SPL13*), four mainly in spikes (*SPL14, SPL16, SPL17* and *SPL18*), and one (*SPL7*) was almost undetectable in all tissues (Figure S1b). In the immature leaves *SPL3, SPL4, SPL13* and *SPL14* showed higher expression than the other *SPL* genes (Figure S1c). Since the objective of this study was to characterize the effect of the wheat *SPL* genes on the regulatory gene network affecting heading time in the leaves, we initially focused on the four miR156-regulated *SPL* genes expressed in mature leaves (*SPL2*, *SPL3, SPL4,* and *SPL13*).

To simplify the genetic analyses, we performed all our experiments in the tetraploid wheat (*Triticum turgidum* ssp. *durum*, genomes AABB) cultivar Kronos. Kronos carries the dwarfing allele *Rht-B1b* (GA-insensitive), the *Vrn-A1* allele for spring growth habit, and the reduced photoperiod sensitivity allele *Ppd-A1a* that confers earlier heading under SD. We first validated the expression of *SPL2*, *SPL3, SPL4,* and *SPL13* in the leaves by qRT-PCR using primers and methods described in Data S2 and Method S2. All four genes showed increased transcript levels in the leaves with plant age (Figure S4a-d) and altered expression in transgenic plants with increased or reduced levels of miR156. *SPL2*, *SPL3, SPL4,* and *SPL13* were significantly down-regulated in the leaves of plants overexpressing miR156 (Figure S4e-h), and up-regulated in plants expressing a target mimicry against miR156 (MIM156, Figure S4i-l, Data S3). *SPL2* transcript levels in leaves were >10-fold lower than those of other *SPL* genes and were significantly upregulated in only two of the five MIM156 transgenic lines (Figure S4l). Based on these results, *SPL2* was excluded from further studies.

Recessive mutations have limited effects in polyploid species, so we targeted the miR156 binding sites of *SPL3*, *SPL4*, and *SPL13* to generate dominant alleles. Using a public database of sequenced EMS-induced mutations in tetraploid wheat Kronos and hexaploid wheat Cadenza (Krasileva *et al*., 2017), we identified mutations within the miR156 target sites for *SPL-A4* in Kronos, and for *SPL-A3* and *SPL-B3* in Cadenza (Figure 1a and b). The mutations from Cadenza were transferred to Kronos by marker assisted backcrossing (see Method S3). We did not find EMS-induced mutations within the miR156 binding site of *SPL13*, so we generated CRISPR-induced mutations within this region for both *SPL-A13* and *SPL-B13* in Kronos (Figure 1c, Method S4). These selected mutations reduce the binding energy of miR156 (Figure 1a-c), resulting in miR156-resistant alleles (henceforth, *rSPL*). All three *rSPL* alleles were expressed at higher levels than their respective wildtype alleles (Data S3, Figure S5).

**Figure 1.**
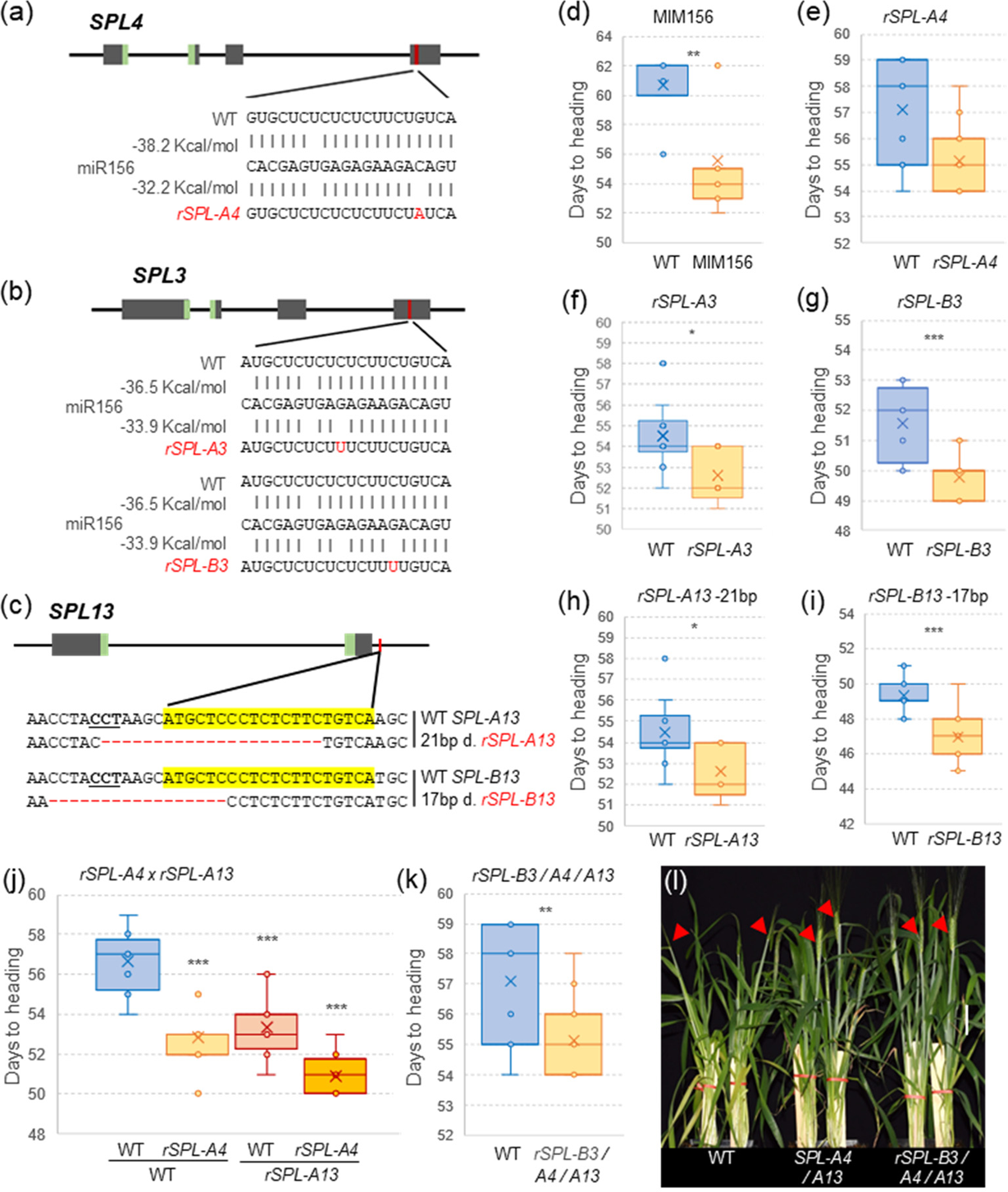
miR156-regulated *SPL3*, *SPL4* and *SPL13* promote flowering time in spring wheat. **(a-c)** Schematic diagrams showing gene structure, conserved SBP domain in green, and miR156 target site in red. Mutations and edits in the sequences are indicated with red letters **(a)** *rSPL-A4* synonymous mutation in Kronos mutant line K2942. **(b)** *rSPL-A3* and *rSPL-B3* mutations in Cadenza (CAD1995 and CAD1033) **(c)** CRISPR induced-deletions in miR156 binding sites of *SPL-A13* (21 bp) and *SPL-B13* (17 bp) located in the 3’ UTR. The miR156 target sequence is highlighted in yellow, and the PAM site is underlined. **(d-k)** Days to heading under long (LD). **(d)** Target mimicry against miR156 (MIM156 n= 15) *vs.* wildtype (WT n= 7) **(e)** *rSPL-A4* (n= 13) *vs.* WT (n= 11). **(f)** *rSPL-A3* (n= 5) *vs.* WT (n= 10). **(g)** *rSPL-B3 vs.* WT (both n= 12). **(h)** *rSPL-A13* (n= 11) *vs.* WT (n= 9). **(i)** *rSPL-B13 vs.* WT (both n= 11). **(j)** *rSPL-A4* and *rSPL-A13* single and double mutants compared to WT (all n= 12). **(k)** Triple resistant lines combining *rSPL-B3 rSPL-A4 rSPL-A13* (n= 15) *vs.* WT (n= 14). **(l)** Comparison between plants carrying wildtype, double *rSPL-A4/A13*, and triple *rSPL-B3/A4/A13* resistant alleles at head emergence. Error bars represent s.e.m. *P* values are from *t*-test (ns= not significant, * *P*< 0.05, ** *P*< 0.01, *** *P*< 0.001). Raw data and statistical analyses are available in Data S4.

In summary, we identified three wheat *SPL* genes targeted by miR156 and expressed in the leaves and generated five dominant miR156-resistant alleles that were expressed at higher levels than their respective wildtype alleles.

### Increased expression of SPL3, SPL4, and SPL13 accelerates heading time and reduces leaf number

We first explored the effect of MIM156 and *SPL* resistant alleles on heading time under long days (LD, Table 1). The MIM156 transgene led to the simultaneous upregulation of multiple *SPL* genes (Figure S4i-k) and accelerated heading by 5.2 days relative to the wildtype (Figure 1d). The individual *rSPL* alleles showed smaller effects on heading time (Figure 1e-i). The synonymous mutation in *rSPL-A4* did not alter the encoded protein but affected the binding energy of the miR156 (Figure 1a) and accelerated heading 2.1 ± 0.5 days (average ± s.e.m. of three experiments) relative to the wildtype (Table 1, Figure 1e).

**Table 1.** Effect of *rSPL* alleles on days to heading (DTH) and leaf number (LN) relative to wildtype alleles in two experiments. Raw data are available in supplemental Data S4, S5, S7, and S8.

| Exp. | Genotype | Photoperiod | Diff. DTH | $P^1$ | Diff. LN | $P^2$ |
| --- | --- | --- | --- | --- | --- | --- |
| 1 | *rSPL-A3* | LD | -1.90 | * | -0.70 | * |
| 3 | *rSPL-A3* | LD | -3.18 | * | -0.82 | * |
| 1 | *rSPL-B3* | LD | -1.83 | *** | -0.43 | * |
| 2 | *rSPL-B3* | LD | -3.40 | *** | 0.00 | ns |
| 3 | *rSPL-B3* | LD | -2.14 | * | -0.29 | * |
| 2 | *rSPL-B3* | SD | -3.03 | * | 0.00 | ns |
| 1 | *rSPL-A4* | LD | -1.94 | ** | -0.50 | * |
| 2 | *rSPL-A4* | LD | -1.33 | ns | -0.33 | ns |
| 3 | *rSPL-A4* | LD | -2.88 | *** | -0.88 | *** |
| 1 | *rSPL-A4* | SD | -6.41 | *** | . | . |
| 2 | *rSPL-A4* | SD | -4.58 | ** | -0.70 | * |
| 1 | *rSPL-A13* | LD | -3.06 | *** | -0.67 | * |
| 2 | *rSPL-A13* | LD | -0.37 | ns | 0.50 | ns |
| 3 | *rSPL-A13* | LD | -2.65 | * | 0.15 | ns |
| 2 | *rSPL-A13* | SD | -4.21 | * | -0.69 | * |
| 1 | *rSPL-B13* | LD | -2.36 | *** | . | . |
| 3 | *rSPL-B13* | LD | -2.89 | ** | -0.25 | ns |
| 1 | *rSPL-A4/A13* | LD | -5.75 | *** | -1.33 | *** |
| 2 | *rSPL-A4/A13* | LD | -2.83 | ** | -0.17 | ns |
| 2 | *rSPL-A4/A13* | SD | -6.43 | ** | -0.86 | ** |
| 1 | *rSPL-B3/A4/A13* | LD | -6.55 | *** | -1.79 | *** |
| 2 | *rSPL-B3/A4/A13* | LD | -3.83 | *** | -1.00 | ** |
| 2 | *rSPL-B3/A4/A13* | SD | -6.34 | ** | -1.00 | *** |
| <sup>1</sup> *t*-Test, <sup>2</sup> Kruskal-Wallis |  |  | Correlation DTH vs LN |  | 0.74 |  |

The mutations in *rSPL-A3* (serine to phenylalanine) and *rSPL-B3* (synonymous mutation) miR156 binding sites were associated with higher *SPL3* transcript levels (Data S3, Figure S5) and similar accelerations in heading time relative to their wildtype sister lines (average 2.5 ± 0.6 days in *rSPL-A3* and 2.5 ± 0.5 days in *rSPL-B3*, Figure 1f-g and Table 1). The similar effects of the *rSPL-A3* and *rSPL-B3* alleles on heading time suggested that the amino acid change in *rSPL-A3* did not affect the function of the encoded protein.

The miR156 binding site of *SPL13* is in the 3’ UTR, so the CRISPR-induced indels in this region in *rSPL-A13* (21-bp deletion) and *rSPL-B13* (17-bp deletion) are not expected to affect the encoded proteins (Figure 1c). In two Cas9-free populations segregating for either *rSPL-A13* or *rSPL-B13*, plants carrying the resistant alleles showed higher *SPL13* transcript levels (Figure S5) and headed 2.0 ± 0.8 days (*rSPL-A13,* Figure 1h) and 2.6 ± 0.3 days (*rSPL-B13*, Figure 1i) earlier than their respective wildtype controls.

We then intercrossed plants carrying the *rSPL-A4* and *rSPL-A13* alleles to study their combined effects. Plants carrying both *rSPL-A4* and *rSPL-A13* alleles headed on average 5.8 days earlier than the wildtype, whereas smaller differences were observed for sister plants carrying *rSPL-A4* (3.8 days) or *rSPL-A13* (3.3 days) alone (Figure 1j). The factorial ANOVA for days to heading (DTH) showed highly significant effects for the individual resistant alleles (*P<* 0.001) and a marginally non-significant interaction (*P=*0.0719, Data S4). These results suggest that the positive effects of the two alleles on DTH are mostly additive. Finally, we combined the three resistant alleles *rSPL-B3*, *rSPL-A4*, and *rSPL-A13*, and observed that plants homozygous for all three mutations headed 6.6 days earlier than the sister lines without any resistant allele (Figure 1k, Data S4). A photograph comparing head emergence in plants carrying the combined resistant alleles and the wildtype is presented in Figure 1l. Similar photographs for the individual *SPL* resistant alleles and their corresponding wildtype sister lines are presented in Figure S6.

To determine which developmental phase was accelerated in the *rSPL* mutants, we measured leaf number (LN), which is fixed at the time of the SAM transition to an IM. Plants carrying the single *rSPL-A3* or *rSPL-B3* alleles showed an average of 0.8 ± 0.1 and 0.24 ± 0.1 fewer leaves than the wildtype, respectively (Table 1, Figure S7a-b). Similar differences were observed in plants carrying homozygous *rSPL-A4* alleles (0.6 ± 0.2 leaves, Figure S7). The effects of *rSPL-A13* and *rSPL-B13* on leaf number were significant in only one of the four experiments conducted under LD (Table 1, Figure S7c).

A factorial ANOVA for LN comparing the individual and combined *rSPL-A4 rSPL-A13* alleles showed significant reductions in LN (*P=* 0.02) for both *rSPL-A4* (0.50 leaves) and *rSPL-A13* (0.67 leaves), and highly significant reductions (*P*< 0.001) for the combined alleles (1.3 leaves, Figure S7d, Table 1). The interaction between the two genes was not significant, indicating additive effects (Data S5). The triple mutant carrying the *rSPL-B3, rSPL-A4* and *rSPL-A13* alleles had on average 1.4 ± 0.4 fewer leaves than the wildtype across two experiments performed under LD (Table 1, Figure S7e).

To further characterize the effect of *SPL13*, we generated transgenic Kronos plants overexpressing the *SPL-A13* coding region fused with a C-terminal HA tag under the maize *UBIQUITIN* promoter (Method S4). This transgene does not include the 3’ UTR with the miR156 binding site and, therefore, is insensitive to miR156 activity (Ubi::rSPL-A13-HA). All five independent transgenic plants showed higher *SPL13* expression levels than the non-transgenic controls in the third leaf, but the differences were significant only for three of them (Figure 2a, Data S6). These same three transgenic plants showed significant increases in *VRN1* expression (Figure 2b), but the differences for *FT1* were not significant (Figure 2c). In agreement with the expression results, all five transgenic plants showed reduced DTH (average 2.1 days, Figure 2d) and LN (average 1 leaf, Figure 2e).

**Figure 2.**
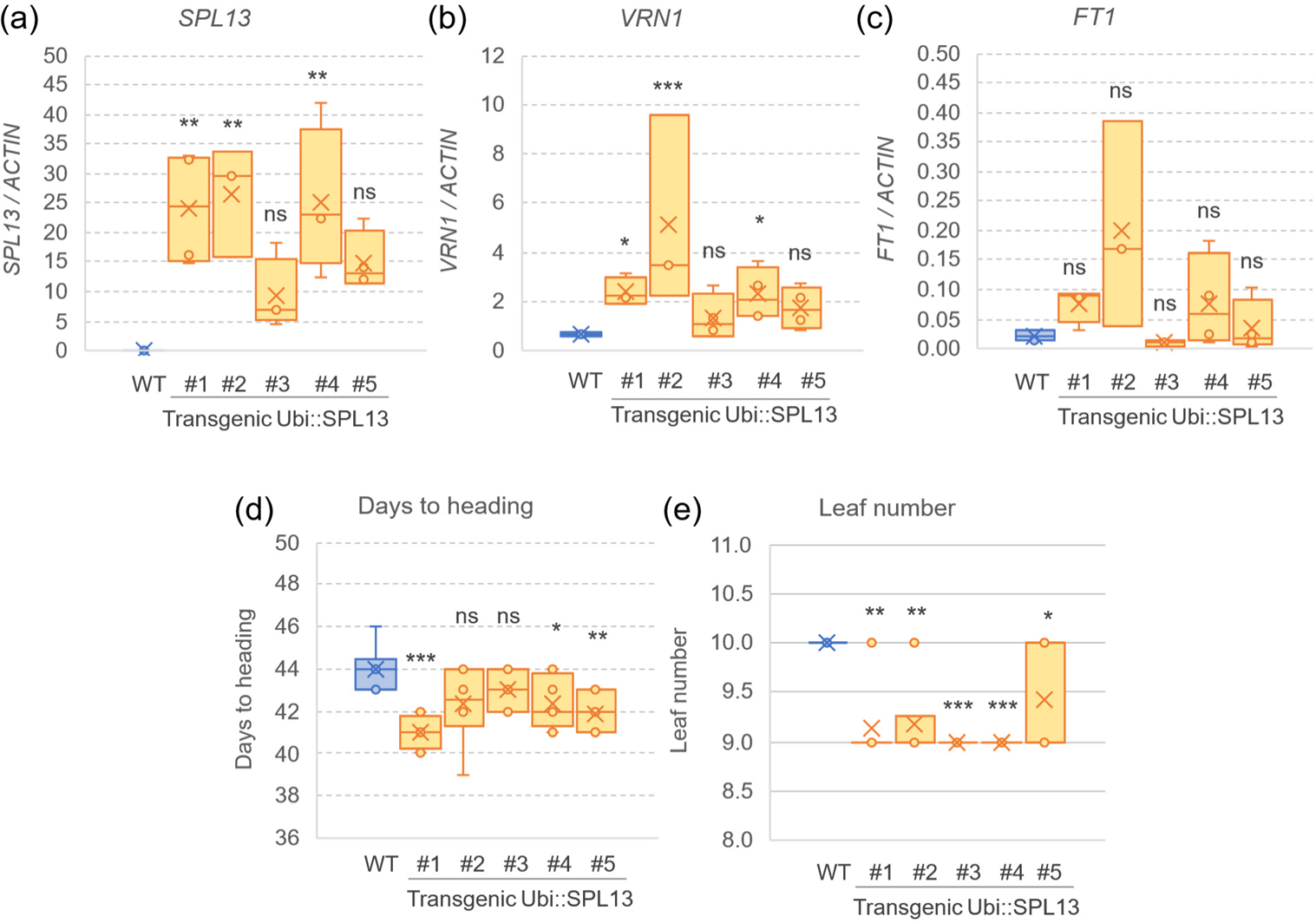
Effect of Ubi::rSPL-A13-HA on gene expression in the leaves and heading time. Transcript levels in the third leaves are expressed relative to *ACTIN* as endogenous control using he Delta Ct method. **(a-c)** Transcript levels of **(a)** *SPL13*, **(b)** *VRN1*, **(c)** *FT1* in the third leaf of Ubi::rSPL-A13-HA and a sister line without the transgene (WT). **(d)** Days to heading and **(e)** Leaf number. Means of the transgenic plants were compared with the wildtype (WT) using Dunnett tests. ns= not significant, *= *P<* 0.05, **= *P<* 0.01, ***= *P<* 0.001. Raw data and statistical analyses are available in Data S6.

In summary, the increased *SPL* expression of the five *rSPL* alleles and Ubi::rSPL-A13 was associated with relatively similar accelerations of heading times and reductions in LN under LD, and the effects of combined alleles were additive.

### rSPL4 and rSPL13 have stronger effect on heading time under SD than under LD

Next, we compared the effect of the *rSPL* alleles on DTH and LN under both LD and SD (Table 1 and Figure S8). For *rSPL-A4*, we performed two experiments, which showed similar results: stronger differences in DTH between genotypes under SD (6.4 d and 4.6 d) than under LD (1.9 and 1.3 d) (Figure S8a-b, Data S7). Factorial ANOVAs including photoperiod and *SPL4* alleles showed highly significant effects for both factors (*P<* 0.0001) and significant interactions in both experiments (*P=* 0.024 and 0.029, respectively, Data S7), confirming the stronger effect of the *rSPL-A4* allele on DTH under SD than under LD.

The *rSPL-A13* allele also showed a stronger effect on heading time under SD, which was supported by a significant genotype x photoperiod interaction in the factorial ANOVA analysis (Figure S8c, Data S7). By contrast, the differences between *rSPL-B3* and its wildtype sister lines were similar under LD and SD (Figure S8d). As expected, the combined double *rSPL-A4/A13* and triple *rSPL-B3/A4/A13* resistant alleles also showed stronger effects on DTH under SD than under LD (Figure S8e-f, Data S7).

Surprisingly, the differences in LN between genotypes were not affected by photoperiod and showed similar values in SD and LD, both for the single and combined mutants (Table 1). These results suggested that the larger differences in DTH under SD relative to LD were associated mainly with changes in the spike differentiation and elongation phase rather than in the transition from the SAM to the reproductive stage. A relatively high correlation between the differences between genotypes in DTH and LN (*R=* 0.74) was observed when single and combined *rSPL* alleles were analyzed together (Table 1).

In summary, the differences between *rSPL* and wildtype alleles were larger under SD than under LD for DTH (for *rSPL-A4* and *rSPL-A13*) but not for LN.

### rSPL alleles upregulate flowering promoting genes and downregulate flowering repressing genes in leaves

To explore the molecular mechanisms underlying the earlier heading of the *rSPL* mutants relative to their sister wildtype lines, we compared the transcript levels of several flowering promoting and repressing genes at different leaves using qRT-PCR. We performed these comparisons in older leaves for plants grown under SD (L5, L7 and L9) than under LD (L3, L5 and L6) to account for the delayed transition to the reproductive phase under SD (Table 2). Despite some variability, an overall analysis of the significant comparisons in Table 2 revealed a consistent upregulation of flowering promoting genes (miR172, *FT1*, *FUL2*, and *VRN1*) and downregulation of flowering repressing genes (*VRN2* and *AP2L1*) associated with the *rSPL* alleles (Data S9). These expression results are consistent with the earlier heading of the plants carrying the *rSPL* alleles.

**Table 2.** Effect of *rSPL* resistant alleles on the expression of main flowering genes under long (LD, 16h light) and short days (SD, 8h light). The yellow highlight indicates higher expression in the resistant allele than in the wildtype, whereas the blue highlight indicates lower expression in the resistant allele than the wildtype. Raw data and statistical tests are available in (Data S9).

| Leaf | Ph.P. | Exp. | Gene | *miR172* | *FT1* | *FUL2* | *VRN1* | *VRN2* | *AP2L1* |
| --- | --- | --- | --- | --- | --- | --- | --- | --- | --- |
| L3 | LD | 1 | *rSPL-A4* | ns | ** | ns | ns | ns | ns |
| L5 | LD | 1 | *rSPL-A4* | * | ** | ** | ns | ns | NA |
| L5 | LD | 2 | *rSPL-A4* | ns | *** | *** | ns | ** | ** |
| L6 | LD | 2 | *rSPL-A4* | * | ns | *** | ** | ** | ns |
| L5 | SD | 2 | *rSPL-A4* | ns | ** | ** | * | *** | ns |
| L7 | SD | 2 | *rSPL-A4* | * | ns | * | * | 0.079 | 0.061 |
| L9 | SD | 2 | *rSPL-A4* | ns | ns | ** | ns | ns | ns |
| L3 | LD | 1 | *rSPL-A3* | ns | ns | ns | ** | * | 0.063 |
| L5 | LD | 1 | *rSPL-A3* | ns | ns | ns | ns | ns | ns |
| L5 | SD | 2 | *rSPL-A3* | ns | ns | ns | ns | ns | 0.097 |
| L7 | SD | 2 | *rSPL-A3* | ns | ns | ns | * | ns | ns |
| L9 | SD | 2 | *rSPL-A3* | 0.055 | ns | ns | ns | ns | ns |
| L3 | LD | 1 | *rSPL-B3* | ns | ns | 0.072 | 0.066 | ns | ns |
| L5 | LD | 2 | *rSPL-B3* | ns | ns | * | ns | 0.053 | ** |
| L6 | LD | 2 | *rSPL-B3* | ns | ns | * | ** | 0.063 | ns |
| L5 | SD | 2 | *rSPL-B3* | ns | ns | ns | * | ns | ns |
| L7 | SD | 2 | *rSPL-B3* | ns | ns | ns | ns | ns | ns |
| L9 | SD | 2 | *rSPL-B3* | ns | ns | ns | ns | ns | ns |
| L3 | LD | 1 | *rSPL-A13* | ns | ns | ns | ns | 0.089 | * |
| L5 | LD | 1 | *rSPL-A13* | ns | ns | ns | ns | ns | ns |
| L5 | LD | 2 | *rSPL-A13* | ns | ns | ns | ns | ns | ns |
| L6 | LD | 2 | *rSPL-A13* | ns | ns | ns | ns | ns | ns |
| L5 | SD | 2 | *rSPL-A13* | ns | ns | ns | ns | ** | ns |
| L7 | SD | 2 | *rSPL-A13* | ns | * | * | ns | * | ns |
| L9 | SD | 2 | *rSPL-A13* | * | * | ns | * | ns | ns |
| L3 | LD | 2 | *rSPL-B13* | ns | ns | ns | ns | ns | ns |
| L5 | LD | 2 | *rSPL-B13* | ns | ns | ** | ns | * | ns |
| L6 | LD | 2 | *rSPL-B13* | ns | * | *** | 0.068 | * | ns |
| L5 | SD | 2 | *rSPL-B13* | ns | * | ns | * | *** | ns |
| L7 | SD | 2 | *rSPL-B13* | ns | ** | ** | ns | ** | ns |
| L9 | SD | 2 | *rSPL-B13* | ** | ns | * | ns | ns | ns |

Comparisons between *rSPL-A4* and their wildtype sister lines showed the highest proportion of significant tests, with significant effects observed in all six genes. A smaller number of assays were significant for the *rSPL3* alleles, and they were primarily concentrated on the *FUL2* and *VRN1* genes. This latter result suggested that these two MADS-box genes may be the initial targets of the SPL3 proteins. Plants carrying the *rSPL13* alleles showed significant difference for all flowering genes, but *AP2L1* was significant for only one comparison in L3. Significant differences for *rSPL13* were concentrated in experiments performed under SD and were more frequent for *rSPL-B13* than for *rSPL-A13* (Table 2), which coincides with the higher expression levels of *rSPL-B13* in leaves (Figure S1b). The positive effect of *rSPL13* on *VRN1* expression was validated in Ubi::rSPL-A13 transgenic plants ectopically expressing *SPL-A13* (Figure 2B).

In summary, the expression results suggest that the *rSPL3*, *rSPL4,* and *rSPL13* alleles accelerated heading time by up-regulating flowering-promoting genes and downregulating flowering-repressing genes in the leaves.

### The miR156/SPL pathway modulates vernalization requirement and heading time in winter wheat

To explore the interactions between the age and vernalization pathways, we introduced the MIM156 cassette into a winter Kronos line carrying a loss-of-function mutation in the *Vrn-A1* allele for spring growth habit (Chen and Dubcovsky, 2012). Plants subjected to different vernalization treatments were synchronized through sequential planting dates and were at a similar developmental stage at the time of removal from the cold treatment.

In the absence of vernalization, the MIM156 winter plants headed 59 days earlier than the control winter Kronos plants without the transgene (Figure 3a). However, this difference was reduced to 34 days after 30 days of vernalization and to 22 days after 42 days of vernalization. A factorial ANOVA for heading time (Data S10) revealed highly significant effects of genotype (*P<* 0.001), vernalization treatment (*P<* 0.001), and a significant interaction (*P*=0.0016, Figure 3a, Data S10) that reflects the increased effects of MIM156 on heading time under suboptimal flowering inductive conditions (Figure 3a).

**Figure 3.**
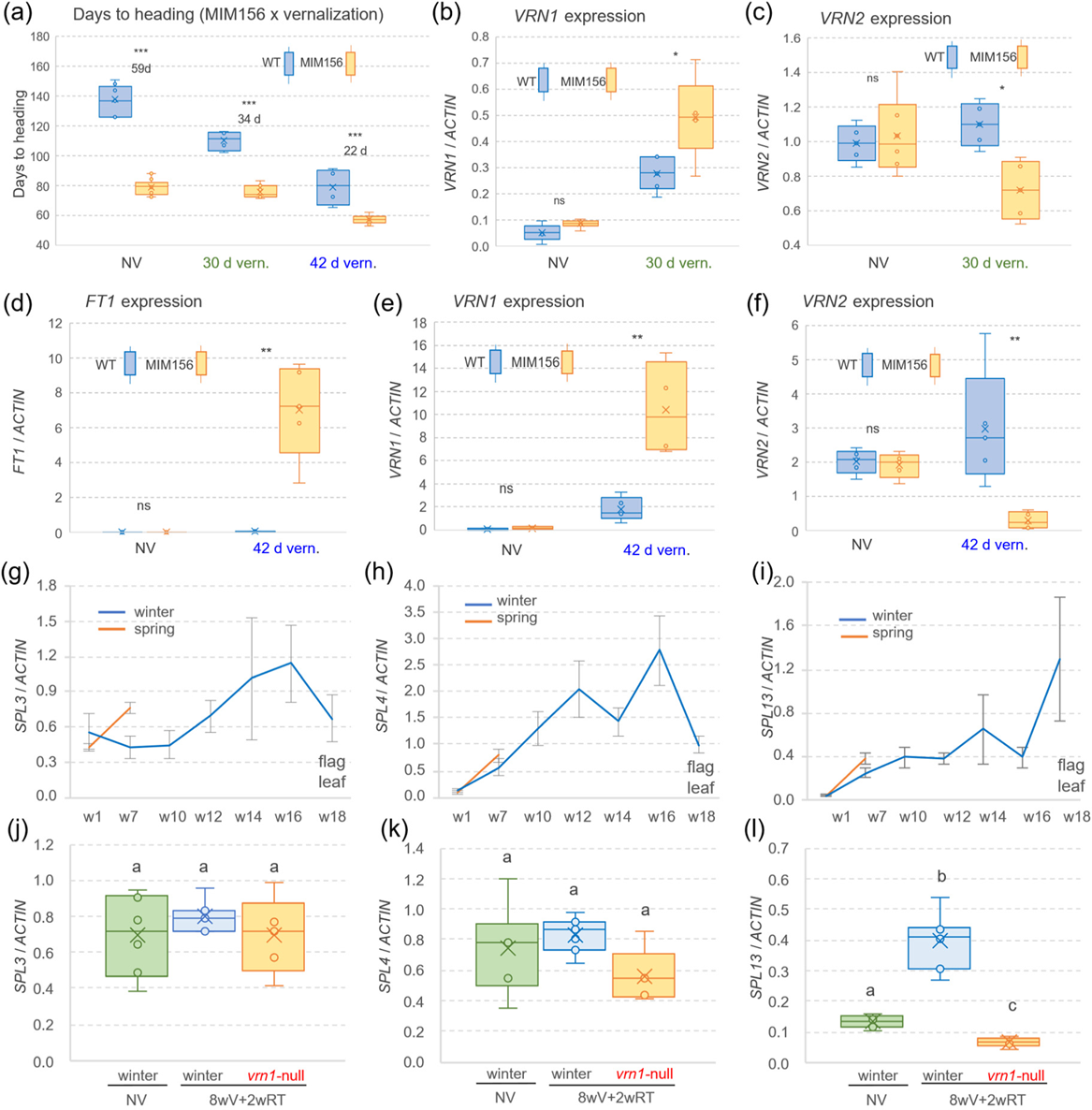
Interactions between the age and the vernalization pathways in wheat leaves. **(a)** Box plot showing days to heading of wildtype and MIM156 plants grown without vernalization (NV) and with 30 or 42 days vernalization. **(b-c)** Expression levels of *VRN1* **(b)** and *VRN2* **(c)** determined in leaves of wildtype (WT) and plants carrying MIM156 without vernalization or after 30 d vernalization followed by two weeks at room temperature (RT). **(d-f)** Expression levels of *FT1* **(d)**, *VRN1* **(e)** and *VRN2* **(f)** in leaves of WT and MIM156 without vernalization and after 42 days vernalization followed by two weeks at RT. **(g-i)** Transcript levels of *SPL3* **(g)**, *SPL4* **(h)** and *SPL13* **(i)** in leaves of spring and winter Kronos plants grown under long days (LD). W= week, L= fully expanded leaf. W1= L1, W5= L7, W10= L10-11, W12= L13-14, W14= L15-16, W16= L18-19, and W18= flag leaves L20-23. **(j-l)** Box plots showing expression levels of *SPL3* **(j)**, *SPL4* **(k)**, and *SPL13* **(l)** in L7 of NV winter Kronos plants, and L7 of winter Kronos and *vrn1*-null mutant plants vernalized for 8 weeks and then moved to RT for 2 weeks (8wV+2wRT). Transcript levels were quantified by qRT-PCR using *ACTIN* as endogenous control. Error bars represent s.e.m. *P* values are from *t*-test ( ns= not significant,*= *P*< 0.05, **= *P*< 0.01,b***=*P*< 0.001). Raw data and statistics in Data S10.

Two weeks after removing plants from the different cold treatments, we collected leaf samples and measured the expression of several flowering genes. *VRN1* transcripts were almost undetectable in the non-vernalized plants, but increased after 30 days of vernalization, with MIM156 plants showing higher expression levels than the control without the transgene (Figure 3b). The *VRN2* transcripts decreased after the partial vernalization treatment in the MIM156 plants but not in the wildtype (Figure 3c). Plants exposed to 42 days vernalization showed higher *FT1* and *VRN1* and lower *VRN2* transcript levels than those treated for 30 days, with significantly larger effects in the MIM156 plants than in the wildtype (Figure 3d-f). These expression profiles were consistent with the earlier heading of the plants carrying the MIM156 transgene relative to the control (Figure 3a) and the significant interaction between age and vernalization pathways in the regulation of heading time.

A previous study showed a sharp decrease in miR156 transcript levels in leaves of both spring and non-vernalized winter Kronos plants during the first five weeks (Debernardi *et al*., 2022). Spring plants headed before the next sampling point, whereas the non-vernalized winter plants continued to produce leaves, with the flag leaf emerging at 18 weeks. During this extended period, the winter plants showed a gradual downregulation of miR156 and upregulation of miR172 in the leaves and a delayed upregulation of *VRN1* and *FT1* until week 16 (Debernardi *et al*., 2022). Using the same RNA samples, we observed that transcript levels of both *SPL4* and *SPL13* increased to similar levels in spring and winter wheat during the first five weeks (Figure 3h-i) and then continued to increase in winter wheat. The initial *SPL3* upregulation was delayed in winter Kronos but then increased progressively with age (Figure 3g). Both *SPL3* and *SPL4* transcript levels peaked at week 16 whereas *SPL13* transcripts increased until week 18 (Figure 3g-i).

In a separate experiment, we compared *SPL* expression levels in RNA samples from the 7^th^ leaf collected from either un-vernalized winter Kronos plants or 8-weeks-vernalized plants after they were returned to room temperature for two weeks. No significant differences were detected for *SPL3* or *SPL4*, but *SPL13* showed significantly higher levels in the vernalized plants carrying a functional *VRN-B1* allele than in the *vrn1*-null mutant plants (Figure 3j-l). This result suggested that *VRN1* or some of its downstream targets contributed to the transcriptional upregulation of *SPL13*, and explained the effect of *VRN1* on the upregulation of miR172 and the downregulation of *AP2L1* reported in the same samples in a previous study (Debernardi *et al*., 2022).

In summary, the effects of miR156 on DTH were stronger in non-vernalized than vernalized plants, and reduced levels of miR156 reduced the vernalization requirement in winter wheat. In the absence of vernalization, *SPL* genes were upregulated with age.

### SPL proteins interact with SQUAMOSA regulatory regions

To test if the positive effects of the *rSPL3*, *rSPL4*, and *rSPL-13* alleles on the transcriptional regulation of *VRN1* and *FUL2* (Table 2) were the result of direct physical interactions, we performed yeast-one-hybrid (Y1H, Method S5, Figure S9) and Electrophoretic Mobility Shift Assays (EMSA, Method S6, Figure S10). We selected three *VRN1* and five *FUL2* regulatory regions carrying putative SPL-binding sites for the Y1H assays (Data S2). However, the *VRN-B1* and *FUL-A2* promoter regions, including a putative SPL-binding motif near the transcription start site, exhibited strong activation by endogenous yeast transcription factors and were excluded from the Y1H assays (Data S2). Instead, we evaluated the *FUL-A2* promoter region (-399 to -374) using EMSA and found clear interactions with all three SPL proteins (Figure S10). The decreased intensity of the shifted DNA-protein band upon the addition of non-labelled DNA (cold-probe) further validated the specificity of the interaction.

The orthologous promoter region of *VRN-A1* including one putative SPL-binding site (-404 to -350, Data S2) was able to interact with SPL4 but not with SPL3 or SPL13 in the Y1H assays (Figure S9a). The region in the first intron of *VRN-B1* (+190 to +244) including a putative SPL-binding site failed to interact with any of the three SPL proteins tested (Figure S9b). This region was not evaluated in the *VRN-A1* homeolog, which has a large deletion in the first intron (Fu *et al*., 2005).

In addition to the *FUL-A2* promoter region near the transcription start site, we evaluated upstream promoter regions in *FUL-A2* (-836 and -666, Figure S9c) and *FUL-B2* (-992 to -826, Figure S9d), both of which contain two putative SPL-binding sites (Data S2). The *FUL-A2* region showed a weak interaction only with SPL4, whereas the similar *FUL-B2* region (93% identity) failed to interact with any of the SPL proteins (Figure S9c-d). One of the putative SPL-binding sites is located closer to the border of the DNA bait in *FUL-B2* than in *FUL-A2* (Data S2) and may have reduced the ability of the *FUL-B2* bait to interact with SPL4. Finally, we evaluated a region in the first intron of *FUL2* that included three putative SPL-binding sites in *FUL-A2* (+2,556 to +2,652) and four in *FUL-B2* (+1,346 to +1,514, Data S2). Both the *FUL-A2* and *FUL-B2* regions showed interactions with SPL3 and SPL4 but not with SPL13 (Figure S9e-f).

In summary, only SPL4 interacted with the *VRN1* promoter region (-404 to -350), whereas all three SPL proteins were able to interact with the homeologous *FUL-A2* promoter region. In addition, SPL3 and SPL4 (but not SPL13) interacted with a region from *FUL2* first intron, suggesting functional differentiation among SPL proteins.

### SPL interactions with DELLA proteins modulate their effects on heading time

We first confirmed that the DELLA-SPL physical interactions previously reported in Arabidopsis (Yu *et al*., 2012, Hyun *et al*., 2016) were conserved in wheat using yeast-two-hybrid (Y2H) assays (Method S7). We used a truncated DELLA protein containing only the C-terminal GRAS domain (RHT1-GRAS), as the full-length protein exhibits autoactivation. We detected positive interactions between RHT1-GRAS and both SPL3 and SPL4, but not with the smaller SPL13 protein (Figure S11a-b). These Y2H interactions were validated by bimolecular fluorescence complementation (BiFC) using rice protoplasts in two independent experiments (Figure S12, Method S8) and by co-IP assays in *N. benthamiana* via *Agrobacterium*-mediated infiltration (Figure S13, Method S9). Taken together, these results confirmed that both SPL3 and SPL4 can interact with DELLA-GRAS *in planta*.

To determine the critical SPL region required for the DELLA-SPL interaction (Figure S11c), we generated two SPL-A3 mutants encoding truncated proteins (Figure S11d). The DELLA-GRAS domain interacted with the truncated SPL-A3-Δ157 protein but not with the shorter SPL-A3-Δ194 (Figure S11e), which is missing 37 amino acids present in SPL-B3-Δ157. These 37-amino acids are well conserved in wheat SPL4 and Arabidopsis AtSPL2, AtSPL10 and AtSPL11, all of which also interact with DELLA in Y2H assays (Yu *et al*., 2012). These results suggested that this 37-amino acid region is important for the SPL-DELLA interaction (Figure S11f).

We then tested if the physical interactions between SPL and DELLA proteins were associated with genetic interactions in a cross between a Kronos line carrying MIM156 and a Kronos line containing a deletion encompassing the GA-insensitive *Rht-B1b* allele, designated hereafter as *rht-B1*-null. This mutant line still has a functional *Rht-A1a* allele and, therefore, is GA-sensitive (Method S3). Analyses of the main effects in the F_2_ progeny showed that plants homozygous for the *rht-B1-*null allele were, on average, 19.4 cm taller than those homozygous for the *Rht-B1b* allele (Figure 4a, Data S11). A smaller effect on plant height was observed in the plants carrying the MIM156 transgene, which were 5.5 cm taller than those without the transgene (Figure 4a, Data S11). By contrast, the effects of MIM156 on heading time (4.4 days earlier, Figure 4b) and leaf number (1.3 fewer leaves, Figure 4c) were stronger than those associated with the *RHT1* alleles (1.7 days earlier and 0.6 fewer leaves).

**Figure 4.**
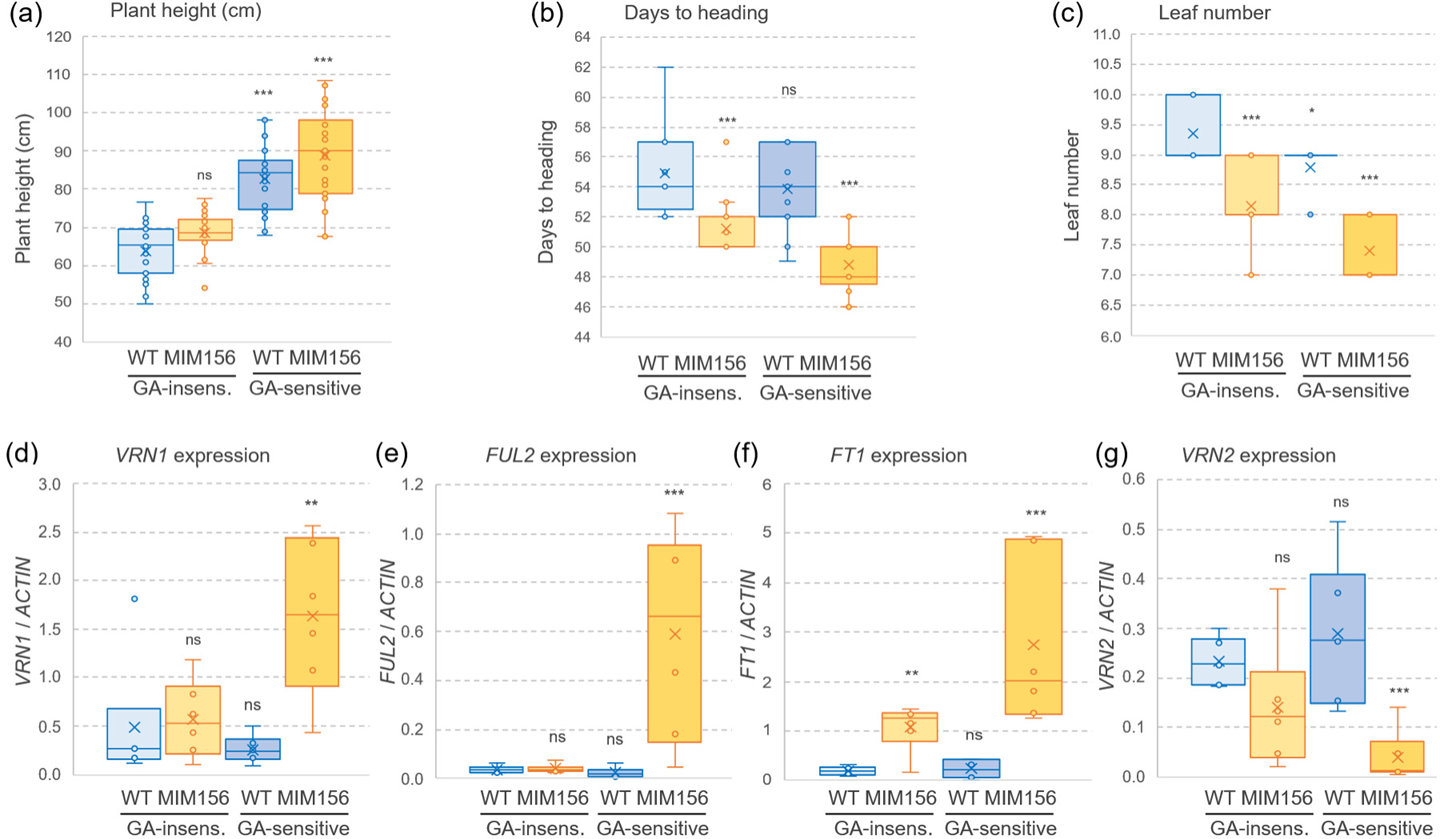
Genetic interactions between the age and GA pathways in wheat leaves. **(a)** Plant height (n= 21-23). **(b)** Days to heading (n= 16-18). **(c)** Leaf number (n= 14-17). Genotypes: wildtype, MIM156, *rht-B1*-null, and double mutants MIM156 *rht-B1*-null. **(d-g)** Expression analysis of *VRN1* **(d)**, *FUL2* **(e)**, *FT1* **(f)** and *VRN2* **(g)** in the third leaves of WT, MIM156, *rht-B1-*null, and double mutants MIM156 *rht-B1*-null. Expression was determined by qRT-PCR using *ACTIN* as endogenous control. Error bars represent s.e.m. ns=not significant, **=*P<* 0.01, and ***=*P<* 0.001 for Dunnett tests vs. wildtype (WT). Raw data and statistics in Data S11.

In addition to the main effects, we evaluated the interactions between MIM156 and *RHT1* (Data S11). For all three traits the effects of MIM156 were stronger in the GA-sensitive background (*rht1-*null, more DELLA) than in the GA-insensitive background (*Rht-B1b*), and the effects of the *RHT1* alleles were stronger in the presence of MIM156 than in its absence (Figure 4 a-c). Although the factorial ANOVAs showed no statistically significant interactions for plant height, DTH, or LN (Data S11), they did reveal significant interactions for the expression levels of flowering genes *VRN1, FUL2*, *FT1*, and *VRN2* in the third leaf (Figure 4d-g, Data S12). Consistent with previous results in Figure S4, plants carrying MIM156 also exhibit a highly significant upregulation of *SPL3*, *SPL4*, *SPL13*, and miR172 (Figure S14, Data S12). The expression results were consistent with the earliest heading of the lines combining MIM156 and *rht-B1-*null alleles and with the interactions for DTH and LN described above. The effects of MIM156 on the expression of these genes were stronger and more significant in the GA-sensitive than in the GA-insensitive backgrounds, and the effects of *RHT1* were stronger and more significant in the presence of MIM156 than in its absence (Figure 4 d-g, Data S12).

In summary, physical interactions were detected between DELLA and both SPL3 and SPL4 proteins, which were paralleled by significant genetic interactions between DELLA and MIM156 for the regulation of flowering genes. Taken together, these interactions suggested that the effects of the GA and miR156 pathways on heading time and leaf number are not fully additive.

### VRN1 and FUL2 interact with DELLA and compete with the SPL-DELLA interaction

Since wheat loss-of-function mutants *vrn1* and *ful2* show reduced plant height (Li *et al*., 2019), we explored the interactions between DELLA and both VRN1 and FUL2. The Y2H assays revealed positive interactions between the DELLA-GRAS domain and both VRN1 and FUL2 proteins, with FUL2 showing stronger interactions than VRN1 (Figure 5a). These interactions were validated by co-IP assays in *N. benthamiana* (Figure 5b, Method S9), confirming that VRN1 and FUL2 can interact with the DELLA-GRAS domain *in planta*.

**Figure 5.**
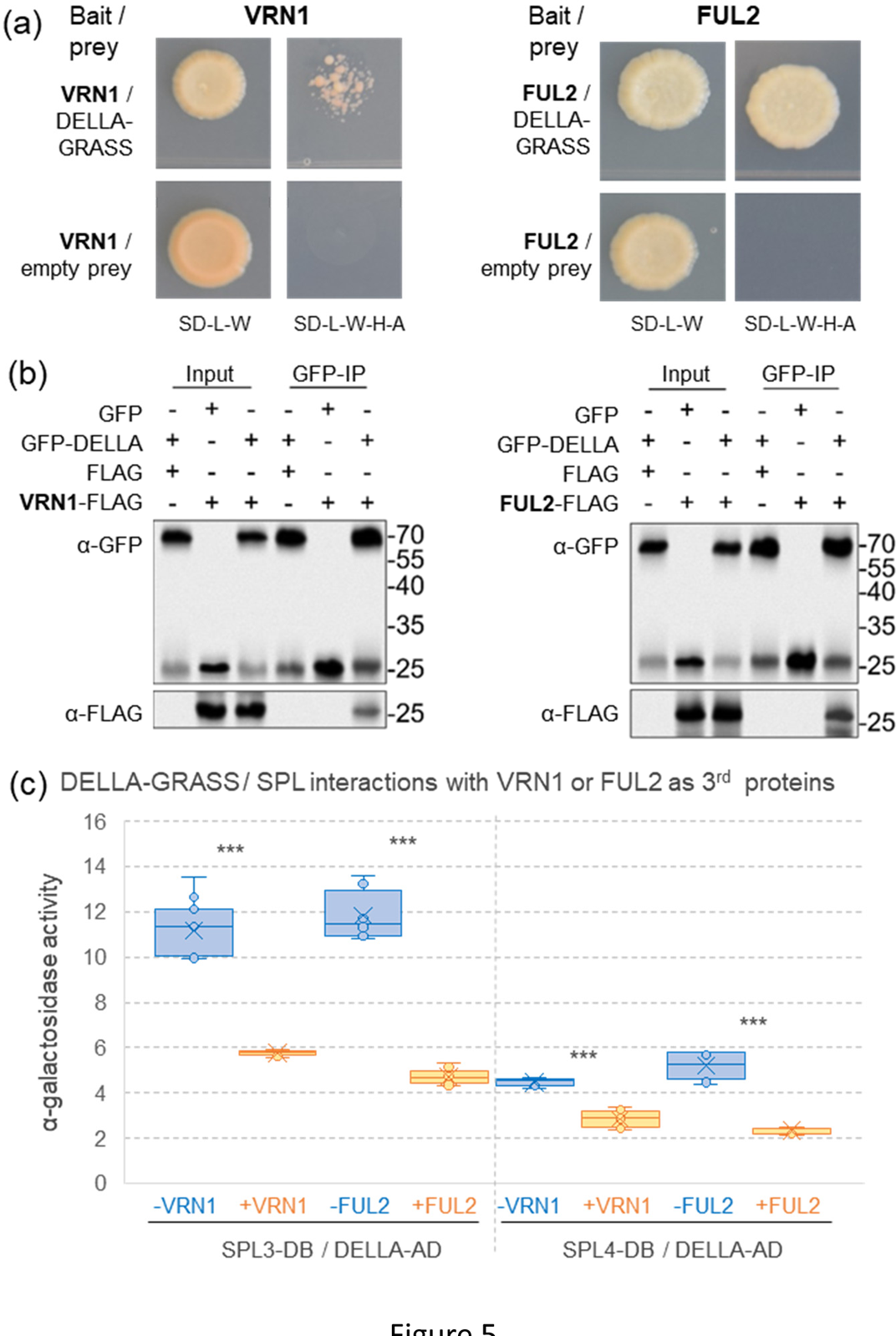
Interactions between DELLA and SQUAMOSA proteins VRN1 and FUL2. **(a)** Y2H assays showing interactions between DELLA and both VRN1 and FUL2 MADS-box proteins and negative controls. **(b)** Confirmation of DELLA - SQUAMOSA interactions by co-IP assays in *N. benthamiana.* Co-IP experiments were performed using GFP-tagged magnetic beads. DELLA-GRASS and the GFP empty vector were detected with the anti-GFP antibody, while VRN1 and FUL2 were detected with the anti-FLAG antibody. The GFP and FLAG empty vectors served as the negative control. **(c)** Alpha-galactosidase activity Y3H assays showing VRN1 and FUL2 competition (as 3^rd^ protein) for the interaction between the DELLA-GRASS domain (activation domain, AD) and SPL proteins SPL3 and SPL4 (DNA binding domain, BD). Lack of autoactivation of the DELLA-GRAS domain and SPL3 and SPL4 proteins is shown in Figure S11a-b. ***= *P<* 0.001. Raw data for the Y3H α-galactosidase activity assays are available in Data S13.

We then used a yeast-three-hybrid system (Y3H, Method S10) to test if the presence of VRN1 or FUL2 expressed as the third protein could interfere with the previously described interaction between DELLA and SPL3 and SPL4 proteins (Figure S11). The quantitative alpha-galactosidase activity assays confirmed the interactions between the DELLA-GRAS domain and both SPL3 and SPL4 proteins and showed a significant decrease in the strength of the DELLA-SPL interactions when either VRN1 or FUL2 was expressed as a third protein (Figure 5c). The presence of FUL2 and VRN1 weakened the DELLA-SPL3 interactions by 60% and 48%, and the DELLA-SPL4 interaction by 55% and 36%, respectively (Figure 5c, Data S13). These results are consistent with the weaker physical interaction between DELLA and VRN1 relative to DELLA-FUL2 (Figure 5a).

In summary, DELLA physically interacted with SQUAMOSA MADS-box proteins VRN1 and FUL2 and these interactions competed with the DELLA-SPL interactions.

### A model for the role of the leaf-expressed SPLs in the regulation of wheat heading time

Figure 6 presents a working model that incorporates the proposed roles of *SPL3*, *SPL4*, and *SPL13*, as well as the interactions among the age, vernalization, photoperiod, and GA pathways in the regulation of wheat heading time. This model focuses on interactions in the leaves that converge on the upregulation of *FT1*.

In winter wheat, the repression of the *VRN1* chromatin is released by vernalization. *VRN1* then promotes *FT1* transcription in the spring by two positive regulatory feedback loops (indicated by circle arrows). The first feedback loop, which has been described in detail in the introduction, involves *VRN1* repressing *VRN2*, the release of *VRN2* repression of *FT1*, and *FT1*’s positive regulation of *VRN1* (Distelfeld *et al*., 2009). The second feedback loop involves the interactions between DELLA and the induced VRN1 and FUL2 proteins, which compete with the repressive DELLA interactions with SPL3 and SPL4 (Figure 5c). The SPL proteins then contribute to the transcriptional upregulation of *VRN1* and *FUL2* by direct binding to their regulatory regions (Table 2, Figure S9 and S10), and indirectly through their positive effect on *FT1* induction (Figure 6). *FT1* is also regulated by the photoperiod pathway through the long-day activation of *PPD1*. The photoperiod pathway is also connected to the endogenous age pathway through the GI upregulation of miR172 (Li *et al*., 2024) (Figure 6).

**Figure 6.**
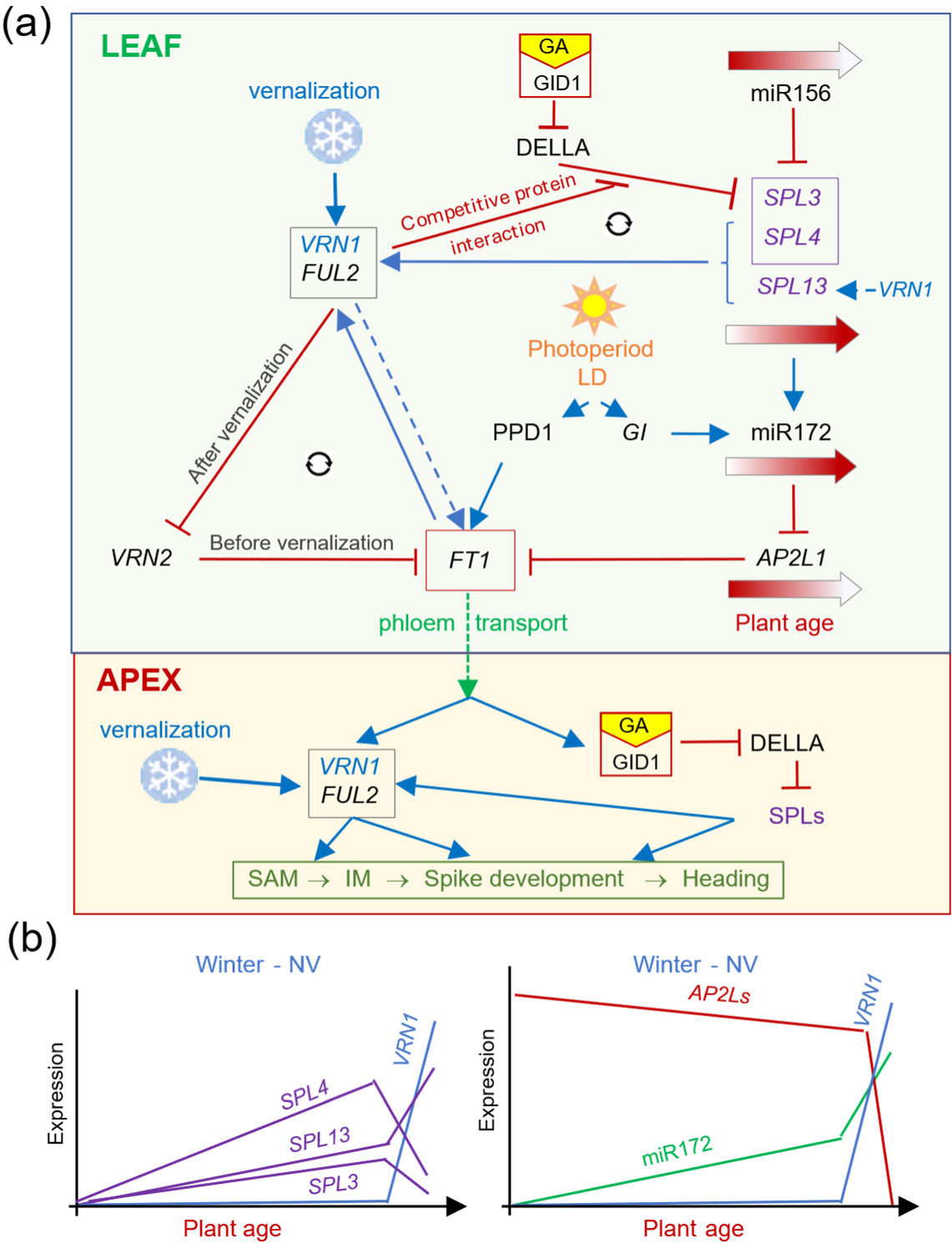
Working model for the role of age pathway in the regulation of wheat heading time. **(a)** The upper panel represents the leaves and the lower panel the shoot apical meristem (SAM) and reproductive tissues. The dotted line from FT1 indicates transport of the protein from leaf to SAM through the phloem. Blue arrows indicate promotion and red lines ending in perpendicular lines indicate repression (dotted blue arrows indicates tentative effects). VRN1 and FUL2 interact physically with the DELLA protein reducing its ability to repress SPL3 and SPL4 protein activity. The three SPL proteins promote the expression of *VRN1*, generating a positive feedback regulatory loop (circular arrows) that accelerates wheat heading time. VRN1, VRN2, FT1 form a separate positive feedback regulatory loop that promotes heading. **(b)** Schematic representation of the expression profiles of *SPL3*, *SPL4*, *SPL13*, *VRN1*, *AP2L* genes and miR172 in the leaves of non-vernalized (NV) winter wheat with very late heading time (18 weeks).

A shorter positive feedback loop has been proposed between *VRN1* and *FT1* based on the reduced expression of *FT1* in the *vrn1* mutants and the binding of VRN1 to the *FT1* promoter in wheat (Tanaka *et al*., 2018) and barley (Deng *et al*., 2015). However, it is difficult to distinguish a direct effect of VRN1 on *FT1* from its indirect effects through *VRN2* and *AP2L1* (Figure 6). A previous study demonstrated that *VRN1* can upregulate *FT1* in a *vrn2-*null mutant background (Shaw *et al*., 2019). However, experiments testing the effect of *VRN1* on heading time in a combined *vrn2-ap2l1*-null background are still pending. To reflect this uncertainty, we connected *VRN1* to *FT1* by a dotted arrow (Figure 6).

The requirement of *VRN1* for flowering is not absolute (Chen and Dubcovsky, 2012) and *vrn1-* null winter wheat Kronos plants eventually head after more than ∼120 days at room temperature and LD conditions (Debernardi *et al*., 2022). The continuous increase of *SPL* transcript levels with age (Figure 3g-I and 6B) and the redundant roles of *FUL2* in spikelet and floral development (Li *et al*., 2019) likely contribute to heading and flowering in the absence of *VRN1*. In summary, this model suggests that the *SPL* genes can regulate *FT1* expression in wheat leaves by both the miR172-*AP2L* and *VRN1-FUL2-VRN2* pathways.

## Discussion

### Distinctive characteristics of the flowering pathways in the temperate cereals

Wheat and other temperate cereals are a monophyletic group characterized by a major expansion and reorganization of their genomes and a common basic chromosome number seven (Kellogg, 2001). These grasses underwent major physiological changes to adapt to cold temperatures, including the development of a vernalization pathway that was not present in the tropical grasses. This pathway is centered on the *VRN1* and *VRN2* genes and involves the epigenetic de-repression of *VRN1* during vernalization (Oliver *et al*., 2009, Liu *et al*., 2024) and its resetting in embryos in the next generation (Niu *et al*., 2024). This pathway is different from the Arabidopsis vernalization pathway, which is centered on the *FLC* and *FRI* genes and involves the epigenetic repression of *FLC* during vernalization (Sung and Amasino, 2005).

The photoperiod pathways in wheat and Arabidopsis also show major differences. While *CO* is the main sensor of day length in Arabidopsis (Andres and Coupland, 2012), it only plays a minor role in wheat, where *PPD1* is a central daylength sensor (Shaw *et al*., 2020). Although the photoperiod pathways of both species converge in the upregulation of *FT1* in the leaves during long days, in winter wheat long days do not induce *FT1* unless *VRN1* is present in the leaves to suppress *FT1* repressors *VRN2* and *AP2L1* (Figure 6). *VRN2* is a grass-specific *FT1* repressor (orthologous to rice *Ghd7*), that is repressed by *VRN1* after vernalization, a pathway that is absent in the tropical grasses (Yan *et al*., 2004b, Dubcovsky *et al*., 2006).

The valuable seasonal information carried by the vernalization-induced VRN1 protein has been integrated into different pathways in the temperate grasses. In winter wheat, for example, the frost tolerance and cold acclimation pathways are less responsive to cold temperatures when *VRN1* is present than when it is absent, resulting in different temperature thresholds in fall and spring (Galiba *et al*., 2009). The induction of *VRN1* also contributes to the down-regulation of the *FT1* transcriptional repressors *VRN2* (Chen and Dubcovsky, 2012) and *AP2L1* (Debernardi *et al*., 2022), thereby generating a more permissive environment for *FT1* induction by the photoperiod pathway (Figure 6). The physical interactions of VRN1 and FUL2 with DELLA represent an additional innovation in the temperate grasses that, to the best of our knowledge, has not been reported before. Our Y3H experiments indicate that these interactions can compete with the DELLA-SPL interactions (Figure 5c), which reduce SPL transcriptional activity in both Arabidopsis (Yu *et al*., 2012) and wheat (Figure 4d-e).

The VRN1/FUL2 – DELLA – SPL physical interactions generate a connection between the endogenous age pathway and the vernalization pathway in wheat, providing a possible mechanistic explanation for the previously reported role of *VRN1* in the repression of *AP2L1* (Debernardi *et al*., 2022). In addition, these competitive physical interactions, together with the ability of SPL3, SPL4 and SPL13 to directly upregulate *VRN1* and *FUL2* transcription, generate a positive feedback loop that reinforces the commitment to flowering (Figure 6). The *SPL* genes can also contribute indirectly to the upregulation of *VRN1* (and possibly *FUL2*) through the induction of miR172, repression of *AP2L1*, and induction of *FT1*, which can bind to the *VRN1* promoter as part of the florigen activation complex and induce *VRN1* transcription (Li *et al*., 2015).

### Conserved functions between wheat and Arabidopsis SPL homologs

Despite the differences between wheat and Arabidopsis in the interactions between the age and other flowering pathways described above, the central endogenous age pathway is relatively well conserved in these two distant species. This conservation includes the downregulation of *miR156* with age and its ability to regulate the *SPL* genes (Figure S4i-k).

Wheat *SPL3* and *SPL4* are related to Arabidopsis *AtSPL2*, *AtSPL10* and *AtSPL11* (Figure S3) and both are expressed at high levels in leaf primordia. However, only wheat *SPL3* and *SPL4* are also expressed at high levels in mature leaves (Figure S1b-c) (Xu *et al*., 2016). Resistant alleles for these *SPL* genes in wheat (Tables 1 and 2) and Arabidopsis (Yao *et al*., 2019) have been associated with the direct transcriptional activation of *SQUAMOSA* MADS-box genes, reduced leaf number, and early flowering. In wheat, SPL4 can bind to a promoter region of *VRN1*, and both SPL3 and SPL4 physically bind to promoter and intron regions of *FUL2* (Figures S9 and S10). In Arabidopsis chromatin immunoprecipitation (ChIP) assays have shown that AtSPL10 binds to the promoters of *AtAP1* and *AtFUL* but not to the first intron (Yao *et al*., 2019).

The wheat *SPL13* gene corresponds to *Arabidopsis AtSPL3*, *AtSPL4*, and *AtSPL5* (Figure S3), and all of them share a small size, absence of conserved domains other than the SBP domain, and a miR156 binding site in the 3’ UTR. The Kronos *rSPL13* allele was associated with a significant acceleration of heading time and altered expression of flowering genes in the leaves, similar to results reported in hexaploid wheat (Gupta *et al*., 2023). The homologous *AtSPL3* gene was expressed in both juvenile and mature leaves but the resistant allele showed no significant acceleration of flowering time (Xu *et al*., 2016) except when expressed under the 35S promoter (Wu and Poethig, 2006, Wang *et al*., 2009). In ChIP assays, AtSPL3 showed preferential binding to regions in the promoter and first intron of *FUL* and *SOC1*, and the promoter of *AP1* (Wang et al., 2009, Yamaguchi et al., 2009). The wheat SPL13 homolog was also able to bind to promoter regions of *FUL2* but not to the region in the first intron including multiple SPL binding sites (Figures S9 and S10).

Overall, we observed a relatively good conservation of the role of the *SPL* genes from these two clades in the acceleration of flowering by the direct upregulation of *SQUAMOSA* genes (Figures S9 and S10), the promotion of *miR172*, and the repression of *AP2L* genes (Table 2).

### Effect of the age pathway under non-inductive conditions

In this study, we show that the effect of the transgenic expression of MIM156 is stronger in non-ernalized winter plants than in spring plants (Figure 1d), with partially vernalized plants showing intermediate effects (Figure 3a). This result is consistent with the increased differences in heading time observed under non-inductive conditions between winter Kronos and its sister lines overexpressing miR172 or carrying combined *ap2l1 ap2l5* mutations (Debernardi *et al*., 2022). These results indicate that genes in the endogenous age pathway can modulate the vernalization response in winter wheat.

We have previously shown that in winter wheat the downregulation of *AP2L1* and the upregulation of *FT1* were decoupled from the age-dependent downregulation of miR156, and that the induction of *VRN1* contributed to the upregulation of miR172, the repression of *AP2L1*, and the activation of *FT1* to promote flowering under LD (Debernardi *et al*., 2022). In this study, we show that these effects can be mediated by SQUAMOSA-DELLA protein interactions competing with the repressive DELLA-SPL3/4 interactions and resulting in more active SPL proteins (Figure 6). In addition, *VRN1* or some of its downstream targets seem to have a positive effect in the transcriptional regulation of *SPL13* (Figure 3l), suggesting that *VRN1* effects on the age pathway are mediated by its transcriptional and post-transcriptional regulation of *SPL* genes.

In non-vernalized winter Kronos plants, the transcript levels of *SPL3*, *SPL4*, and *SPL13* gradually increased, reaching levels higher than those observed in heading spring wheat plants, which likely contributed to the ability of winter Kronos to head after a long time at non-vernalizing temperatures (Figure 3g-i). This hypothesis was supported by the accelerated heading of non-vernalized winter plants overexpressing MIM156, which showed a significant transcriptional upregulation of *SPL3*, *SPL4*, and *SPL13* (Figure S4i-k). These results support the idea that *SPL* genes can accelerate heading time of non-vernalized winter wheat.

This study also showed that *SPL* alleles can accelerate heading time under non-inductive SD, and that the differences between plants carrying wildtype and resistant alleles *rSPL4* or *rSPL13* were larger under SD than under LD (Table 1 and Figure S8). The stronger effect of *rSPL13* on heading time under SD was also supported by field experiments in hexaploid wheat, in which *rSPL13* accelerated heading time by 11 days under fall-planting and by 5 days under spring planting (Gupta *et al*., 2023). Stronger differences in heading time were also reported between Ubi::miR172 and MIM172 transgenic Kronos plants under SD (40 days) than under LD (12 days) (Debernardi *et al*., 2022).

Taken together, the vernalization and photoperiod studies show that wheat genes of the age-dependent pathway have larger effects on heading time under non-inductive conditions than under inductive conditions. These results suggest that the age pathway may serve as a backup system to ensure wheat reproductive development under sub-optimal inductive environments, as previously suggested in Arabidopsis (Wang *et al*., 2009).

### Interactions between the age and the GA pathways modulate wheat heading time

In Arabidopsis, GA signals that promote flowering under non-inductive SD are mediated by the SPL-SOC1 module (Jung *et al*., 2012). GA degrades the DELLA proteins, disrupting the physical interactions with AtSPL9 and AtSPL15 (homologs of wheat SPL14 and SPL17) and AtSPL2/10/11 (homologs of wheat SPL3 and SPL4), restoring SPL activity (Yu *et al*., 2012, Hyun *et al*., 2016). In addition, post-transcriptional and post-translational regulation of AtSPL15 integrate the age and GA pathways at the SAM to promote Arabidopsis flowering under non-inductive conditions (Hyun *et al*., 2016).

Previous studies have also revealed important roles of GA on reproductive development in diploid wheat *Triticum monococcum.* Diploid wheat accessions carrying the photoperiod sensitive allele *Ppd1b* failed to head under SD and did not respond to the addition of exogenous GA. However, accessions carrying the *Vrn1g* or *Vrn1f* alleles with promoter mutations in a CArG box close to the transcriptional start site showed increased transcript levels of *Vrn1* under SD (Dubcovsky *et al*., 2006) and responded to GA applications with accelerated spike development and stem elongation (Pearce *et al*., 2013). These effects were abolished by the addition of the GA inhibitor paclobutrazol, documenting the importance of GA in the promotion of spike development and stem elongation in wheat (Pearce *et al*., 2013). Under natural conditions, the longer days of spring induce *FT1*, triggering the upregulation of *VRN1* and GA biosynthetic genes, which together promote spike development and stem elongation (Pearce *et al*., 2013).

The physical interactions between DELLA and SPL3/4 proteins (Figure S12 and S13) revealed a conserved link between the GA pathway and the endogenous age pathway. The connection between these two pathways was also supported by significant genetic interactions between MIM156 and *Rht1* alleles for the expression levels of flowering genes *VRN1*, *FUL2*, *FT1* and *VRN2* (Data S12). These significant interactions revealed stronger effects of MIM156 on the expression of flowering genes and on heading time in the presence of the GA-sensitive allele, which were likely mediated by reduced DELLA levels and increased SPL activity. These interactions also resulted in stronger effects of the *RHT1* alleles on gene expression and heading time in the presence of MIM156. As a result of these interactions, plants combining MIM156 and GA-sensitive alleles had the highest levels of flowering promoting genes and the lowest levels of flowering repressing genes (Data S12), a result that is consistent with their earliest heading and fewest leaves (Data S11).

We speculate that the elevated levels of DELLA proteins in wheat cultivars carrying GA-insensitive *Rht1b* alleles may exacerbate the repressive effect of DELLA on SPL proteins, and that this effect can be compensated by the induction of VRN1 and FUL2 proteins and their interactions with DELLA. The presence of the SQUAMOSA proteins may reduce the ability of DELLA to repress the SPL proteins and ameliorate DELLA’s negative effect on heading time. *VRN1* contribution to the transcriptional upregulation of *SPL13* (Figure 3l), which does not physically interact with DELLA, may provide an alternative mechanism to offset the accumulation of DELLA in semidwarf wheats carrying the GA-insensitive *Rht1b* alleles.

In summary, we propose that the competitive interactions between DELLA, SQUAMOSA and SPL proteins interconnect the GA, age, and vernalization pathways, and that they can ameliorate the negative effects of the increased DELLA accumulation on heading time in wheats carrying GA-insensitive *Rht1b* alleles.

### Conclusion and practical applications

Wheat is a young polyploid species with high levels of gene redundancy that mask the effects of most recessive mutations (Uauy *et al*., 2017). Therefore, the dominant *rSPL* mutations are particularly attractive for fine tuning heading time in polyploid wheats. In spring wheats like Kronos, the individual *rSPL3*, *rSPL4*, and *rSPL13* alleles accelerate heading time by one to three days, but different alleles can be combined to accelerate heading time by four to six days. In winter wheat and fall-planted spring wheat, which face a longer period of non-inductive conditions, the differences are expected to be one or two days larger than in spring planted wheats. An interesting application of the *rSPL* alleles is to reduce the vernalization requirement in regions where winters become milder due to climate change.

The non-transgenic nature of the *rSPL3* and *rSPL4* mutants makes them particularly attractive for wheat breeding applications because they are not constrained by governmental regulations or CRISPR intellectual property protections that could limit or delay their deployment. To facilitate the adoption of these genetic resources, we deposited the five individual resistant alleles in GRIN Global and made them immediately available without any restrictions (see accession numbers under Data availability).

## Acknowledgements

We thank Xiaoqin Zhang for the transfer of the *rSPL3* mutant alleles from Cadenza to Kronos and Daniel Woods for his help with the phylogenetic tree.

## Supporting information

### Supplemental figures

**Figure S1.** miR156 binding sites in nine wheat SPL genes and their expression profiles.

**Figure S2.** Alignment of SBP domains for Arabidopsis, rice and wheat SPL proteins.

**Figure S3.** Phylogenetic tree of SPL proteins based on the SBP domain alignment.

**Figure S4.** Regulation of SPL3, SPL4, SPL13 and SPL2 expression by miR156.

**Figure S5.** Expression of wildtype and resistant alleles rSPL3, rSPL4, and rSPL13.

**Figure S6.** Pictures of wildtype and rSPL3, rSPL4, and rSPL13 mutants at heading time.

**Figure S7.** Effect of rSPL3, rSPL4, rSPL13 on leaf number.

**Figure S8.** Effect of rSPL3, rSPL4, rSPL13 on heading time under LD and SD.

**Figure S9.** Yeast-one-hybrid (Y1H) assays.

**Figure S10.** Electrophoretic Mobility Shift Assay (EMSA).

**Figure S11.** Interactions between DELLA and SPL proteins.

**Figure S12.** Bi-molecular fluorescent complementation (BiFC or split YFP).

**Figure S13.** CoIP experiments showing interactions between DELLA, SPL3 and SPL4 proteins.

**Figure S14.** Effects of miR156 and Rht-B1 on gene expression.

### Supplemental methods

Method S1. Phylogenetic analysis of SPL family.

Method S2. Expression studies by qRT-PCR.

Method S3. Plant materials, mutants, and growth conditions.

Method S4. Transgenic plants and wheat transformation.

Method S5. Yeast-one-hybrid (Y1H) assays.

Method S6. Electrophoretic Mobility Shift Assays (EMSA).

Method S7. Yeast-two-hybrid (Y2H) assays.

Method S8. Bimolecular fluorescence complementation (BiFC).

Method S9. CoIP assays

Method S10. Yeast-three-hybrid (Y3H) assays.

Method S11. Statistical analyses.

### Supplemental data (S1-S13)

Data S1. Wheat *SPL* nomenclature, gene IDs, and homology with rice *SPLs*.

Data S2. Primers used in this study.

Data S3. Effect of age, miR156, and mutations in miR156 binding site on *SPLs* expression.

Data S4. Effect of *rSPL* alleles on heading time under long days.

Data S5. Effect of single and combined *rSPL3*, *rSPL4*, and *rSPL13* alleles on LN under LD.

Data S6. Effect of Ubi::rSPL-A13HA on gene expression, heading time, and leaf number.

Data S7. Effect of *rSPL* alleles on heading time under long and short-day conditions.

Data S8. Leaf number in plants with single and combined *rSPL* alleles.

Data S9. Effect of *rSPL* alleles on the expression of flowering genes.

Data S10. Effect of age and vernalization on heading time and expression of flowering genes.

Data S11. Interactions between MIM156 and DELLA for plant height, DTH and LN.

Data S12. Interactions between MIM156 and DELLA for expression of flowering genes.

Data S13. Yeast three-hybrid (Y3H) assays.

