## Supplemental Methods for "MicroRNA156 and its targeted *SPL* genes interact with the photoperiod, vernalization, and gibberellin pathways to regulate wheat heading time"

### Supplementary Methods

#### *Method S1. Phylogenetic analysis of the SPL family*

We identified 18 *SPL* genes in the wheat A genome using sequences from rice and *Brachypodium* and combined their encoded proteins with rice and Arabidopsis SPL proteins for a phylogenetic analysis. We used only the conserved SBP domain from these proteins, which includes 53 amino acids. We used the Maximum Likelihood method and Le-Gascuel model (Le & Gascuel, 2008) to generate the final phylogenetic tree. Initial tree(s) for the heuristic search were obtained by applying the Neighbor-Join and BioNJ algorithms to a matrix of pairwise distances and then selecting the topology with the superior log likelihood value. The tree was drawn to scale, with branch lengths measured in the number of substitutions per site. Evolutionary analyses were conducted in MEGA X (Kumar *et al.*, 2018).

#### *Method S2. Expression studies by qRT-PCR*

RNA samples were extracted using the Spectrum Plant Total RNA Kit (Sigma-Aldrich). Protocol A was followed, which allowed purification of total RNA including small RNA molecules. Total RNA was incubated with RQ1 DNase (Promega) for DNA digestion. cDNA synthesis was carried out using SuperScript II Reverse Transcriptase (Invitrogen) from 1 μg of total RNA. Mature miR172 and miR156 levels were detected by stem-loop qRT-PCR as described previously (Chen *et al.*, 2005). Reverse primer for the *Small nucleolar RNA 101* (*SnoR101*) was also included in the reverse transcription to normalize miRNA in qRT-PCR. Quantitative PCR was performed using SYBR Green and a 7500 Fast Real-Time PCR system (Applied Biosystems). *ACTIN* was used as an endogenous control for mRNAs, and *SnoR101* for miRNAs. Primers for the different genes tested are listed in Data S2. Samples were collected from fully extended leaves 3, 5 and 6 under LD and leaves 5, 7 and 9 under SD.

#### *Method S3. Plant materials, mutants, and growth conditions*

The tetraploid wheat variety Kronos (*Triticum turgidum* ssp. *durum* L., 2n = 28, genomes AABB) used in this study carries the *Ppd-A1a* allele (promoter deletion), which confers early heading under short days and is referred to in the literature as “photoperiod insensitive”. Kronos also carries the *Vrn-A1c* allele (intron 1 deletion), which confers a spring growth habit. Finally,

Kronos carries the dominant *Rht-B1b* allele, which encodes an N-truncated DELLA protein that prevents GID1 binding and confers a GA-insensitive semi-dwarf phenotype.

The winter-Kronos near isogenic line (K2268) carries a truncation mutation in the *VRN-A1* genes, and the functional *vrn-B1* gene results in a strong vernalization requirement (Chen & Dubcovsky, 2012). The tall-Kronos near isogenic line (K3822) is the result of 1.9 Mb deletion encompassing the whole *Rht-B1b* gene (Mo *et al.*, 2018), that is designated hereafter as *rht-B1*-null. This deletion still carries the ancestral *Rht-A1a* allele in the A genome, which encodes a wildtype DELLA protein that can be degraded by GA-signaling.

A sequenced Kronos mutant population generated by ethyl methane sulfonate (EMS) (Krasileva *et al.*, 2017) was used to screen for *rSPL* mutants. The synonymous mutation (K2942) detected in the miR156 target site in *SPL-A4* was confirmed in $M_4$ grains using genome specific primers described in Data S2 (Figure 1a). The resulting miR156-resistant allele (*rSPL-A4*) was crossed to the wildtype Kronos and then backcrossed three times to reduce background mutations. Homozygous $BC_3F_2$ sister lines with and without the mutation were used in the experiments described in this study.

For *SPL3*, we did not find good mutations in Kronos, so we selected two EMS mutations in the $M_4$ grains of hexaploid wheat Cadenza: CAD1995 for *rSPL-A3* and CAD1033 for *rSPL-B3*. The mutation in *rSPL-A3* results in an amino acid change from serine to phenylalanine (S389F in CS RefSeq v1.1 TraesCS6A02G110100 and S393F in Kronos v1.0 TrturKRN6A01G018670.2), whereas the mutation in *rSPL-B3* is a synonymous mutation. The mutations were confirmed using genome specific primers described in Data S2 and transferred to Kronos by backcrossing. Crosses between Kronos and Cadenza are not viable due to hybrid necrosis, so we first crossed CAD1033 and CAD1995 with a bridge $F_2$ line derived from the cross between the hexaploid cultivar Insignia and Kronos and then backcrossed each of the $F_1$s two more times with Kronos.

We did not find mutants in the miR156-target site of *SPL13*, so we generated CRISPR mutations for the two *SPL13* homeologs. Since the miR156 target site in *SPL13* is in the 3' UTR, small deletions do not disrupt the encoded protein. Among the five $T_0$ Kronos plants we identified two mutant lines with large deletions disrupting the miR156 binding site: *rSPL-A13* showed a 21-bp deletion in the A genome homeolog and *rSPL-B13* showed a 17-bp deletion in the B genome homeolog.

For all experiments, seeds were germinated in petri dishes for 2-4 days at 4°C. After the first leaf emerged, the seedlings were transplanted to one-gallon pots in growth chambers adjusted to 16 h of light (22°C) and 8 h of darkness (18°C) (long days) or 8 h of light (22°C) and 16 h of darkness (18°C) (short days). The intensity of the sodium halide lights measured at plant height was (~260-300 µM $m^{-2}$ $s^{-1}$). Heading time was recorded from germination until the emergency of half of the main spike from the flag leaf. Plants at the two-leaf stage were vernalized in one-gallon pots in a Conviron PGR15 chamber with temperatures averaging 5°C and long day (16 h of light/8 h of darkness). The intensity of the light was adjusted to 230 uM $m^{-2}$ $s^{-1}$.

***Method S4. Transgenic plants and wheat transformation***

We employed the CRISPR/Cas9 system to create *rSPL13* mutants. We designed a gRNA based on the miR156 target site (Figure 1c) and cloned it into the highly efficient editing vector JD633, which includes the *Cas9* gene and a GRF4-GIF1 chimera that increases wheat regeneration efficiency for *Agrobacterium*-mediated transformation. Five independent $T_0$ transgenic Kronos plants were obtained from the UC Davis Plant Transformation Facility and screened for mutations in *SPL13* genes in the miR156 targeted sites using mid-seq (https://dnacore.mgh.harvard.edu/new-cgi-bin/site/pages/index.jsp). For the next-generation sequencing screen, we used primers SPL13-F1 and SPL13-R1 (Data S2) that can amplify both genomes. Three $T_0$ plants showed editing for A or B genome and their $T_1$ progeny were evaluated to confirm editing. The $T_1$ plant with confirmed editing was crossed with Kronos, and in the progeny, we selected plants homozygous for the mutation and without the transformation vector using primers published previously (Zhang *et al.*, 2025).

Transgenic Kronos plants overexpressing *SPL13* were generated using the Japan Tobacco (JT) vector pLC41 (hygromycin resistance). The coding region of the *SPL13* gene from tetraploid wheat Kronos without the stop codon was cloned in a pDONRTM/Zeo vector and transferred to the pLC41 vector via Gateway LR reaction. The resulting construct has the *SPL13* coding region driven by the maize UBIQUITIN promoter with a C-terminal 3×HA tag and a NOS terminator. Agrobacterium strain EHA105 was used to infect Kronos immature embryos. Transgenic Kronos plants were generated at the UC Davis Plant Transformation Facility (http://ucdptf.ucdavis.edu/)

and positive plants were selected using hygromycin. Transgene insertion was validated by DNA extraction and PCR in $T_0$ using the genome specific primers described in Data S2.

***Method S5. Yeast-one-hybrid assays (YIH)***

The Matchmaker Gold One-Hybrid System (Takara Bio, CA, USA) was used for Y1H assays. Complementary oligos corresponding to three *VRN1* promoter segments and five *FUL2* regulatory regions, each containing one or more putative SPL binding sites (GTACG and TGTACT), were cloned into the *Hind*III-*Xho*I-linearized pAbAi vector (Clontech, CA, USA) upstream of the AUR1-C reporter gene (conferring Aureobasidin A resistance) to generate the bait vectors. After sequence confirmation, each bait vector containing putative SPL binding sites was linearized with either *BstB*I or *Bbs*I restriction enzyme and integrated into the genome of Y1H Gold yeast to generate the bait strain. The minimal inhibitory concentration of Aureobasidin A (AbA) was next determined for each bait strain. Two of the eight strains (#2, containing a *VRN1* promoter segment, and #5, containing a *FUL2* promoter fragment, Data S2) exhibited strong activation by endogenous yeast transcription factors and were excluded from the Y1H assays. For the remaining two *VRN1* bait strains, an AbA concentration of 1000 ng/ml was sufficient to significantly suppress basal AUR1-C expression. For all four *FUL2* bait strains, 200 ng/ml AbA was sufficient. The Kronos coding sequences of SPL-B3, SPL-A4, and SPL-A13 (including the stop codon) were cloned into the pGADT7 vector (Clontech, CA, USA) to generate the prey constructs. Each prey construct was transformed into the individual bait strains following the manufacturer's protocol, and the empty pGADT7 vector was used as a negative control. After transformation, yeast cells transformed with either SPL-pGADT7 constructs or the empty pGADT7 vector were plated on minimal synthetic defined (SD) medium lacking Leucine (SD-Leu) and on SD-Leu supplemented with 200 ng/ml or 1000 ng/ml AbA and then incubated at 30°C for four days to evaluate DNA-protein interactions.

***Method S6. Electrophoretic Mobility Shift Assays (EMSA)***

*Protein expression and purification.* The coding sequences for SPL3**,** SPL4 and SPL13 (sequences shown below) were cloned into the pGEX-6P-1 vector to generate N-terminal GST-tagged fusion proteins. The constructed plasmids were transformed into *Escherichia coli* BL21 (DE3) competent cells. Positive transformants were selected by colony PCR and verified by DNA sequencing. For protein expression, a single verified colony was inoculated into LB medium and cultured overnight. The pre-culture was diluted 1:10 into 1 L of fresh LB medium and grown at 37°C until the $OD_{600}$ reached 0.5-0.6. Protein expression was induced by adding 0.5 mM isopropyl β-d-1-thiogalactopyranoside (IPTG), followed by incubation at 18°C for 12 hours. Cells were then harvested by centrifugation and lysed. The GST-tagged proteins were purified from the clarified lysate using Glutathione Sepharose affinity chromatography. Briefly, the lysate was loaded onto a column packed with 5 mL of Glutathione Sepharose beads pre-equilibrated with equilibration buffer (140 mM NaCl, 2.7 mM KCl, 10 mM $Na_2HPO_4$, 1.8 mM $KH_2PO_4$, pH 7.4). The column was washed extensively with the same buffer, and the bound proteins were eluted with elution buffer (equilibration buffer supplemented with 20 mM reduced glutathione). The purified proteins were analyzed by SDS-PAGE.

```
>TaSPL13
MDRKDKSRKSSSAASMAALAAAAAAGDVARADGMSGEEDQKLKLVNVPVVSVGGSSSSAAAVAVRRGSGAAGAVATGAAGAGGPSCQAERCGADLSE
AKRYHRRHKVCEAHSKAAVVVVAGLRQRFCQQCSRFHELLEFDDQKRSCRRRLAGHNERRRKSSAEANGGDGCRHADQDGRSHPGNPPLNHFQIR*
```

>*TaSPL13* (with codon optimization)

```
ATGGATAGGAAGGACAAATCAAGAAAAAGTTCCTCTGCAGCTAGCATGGCAGCGCTGGCAGCGGCTGCGGCGGCGGGCGACGTTGCCCGTGCAGACG
GCATGAGCGGCGAGGAGGATCAGAAGCTGAAATTAGTTAATGTTCCGGTTGTTAGCGTTGGTGGCTCGTCCTCGTCCGCCGCTGCGGTCGCTGTGCG
TCGCGGTAGCGGTGCGGCTGGCGCGGTGGCCACCGGTGCGGCGGGTGCTGGCGGTCCGAGCTGCCAGGCAGAGCGTTGTGGTGCCGACCTCAGCGAA
GCCAAACGTTACCATCGTCGCCACAAGGTGTGCGAAGCGCACAGCAAAGCAGCGGTGGTGGTGGTCGCCGGGCTGCGTCAGCGTTTCTGCCAACAAT
GTAGCCGATTCCACGAACTGTTGGAATTTGATGATCAGAAGCGCTCCTGCAGACGCCGCCTGGCAGGCCACAACGAGCGCCGTCGTAAAAGCTCTGC
GGAGGCGAACGGTGGTGACGGCTGCCGTCACGCGGACCAAGATGGCCGTTCACATCCGGGTAACCCGCCATTGAATCATTTTCAGATCCGTTAA
```

>*TaSPL3* (with disrupted miR156 binding site)

```
ATGGGCTCTTTTGGGATGGAGTGGAACCAGAGGAGCTCGGTGCTGTGGGACTGGGAGAATTTCCCGCCGATAGGCAATAACCCGAAGAACGCGATGC
AGGCTGATCCAAGATTCGCCGGCGTTGCGGCTACCATGGGGAATGAACCGCTCCATTCTTCTGGCGGCAGCGGCACCTTCTCTTCCAGCTCGGAGAT
GGGGTATGGCTCTTCCAAGAGCTCCATGTCCGCGTCGATCGATTCTTCGTCCAGGGCTGGCAACAACATGGAGTTCAGATTTGCGCCTGTCAAAAAC
CCTGACAGGAACACGAGCAAAAACCCTGAGTTGGGTAAAGTTGACAACACGAGGACTGGAACATCTCCGTCGTCTGTGGTGGCAGTGAGCAGTGGAG
AGCCGGTGATCGGCCTGAAGCTTGGCAAGAGAACTTACTTCGAGGATGTCTGTGGAGGGCAGAGTGTCAAGAGCTTGCCATCGGGTGCTGCGAGCGC
GCCAAACAAATCTCCTGCTTTGGGCAAGAAGGCAAAGGCGGAACAACAGAAGCCACATAACTCATACTGTCAGGTTGAAGGCTGCAAAGCCGATCTC
TCTTCTGTTAAAGATTACCATAGGAAGCACAGGGTCTGTGAACTTCACTCTAAGGCTCCCAAAGTTGTTGTCGCTGGTCTGGAGCGACGCTTTTGCC
AACAGTGCAGCCGGTTTCATGCTTTAGCTGAGTTTGACCAGAAAAAGCGAAGCTGCCGTAGGCGTCTCAATGATCATAATTCCCGCAGGCGCAAGCC
ACAGCCAGAAGCAATTTCTTTCAGTTCATCAAGGATGTCTACGATGTTTTATGATGCAAGGCAACAGCCTAATTTCCTATTTGGTCAGGCTCCTTAT
GTTCAAATGAGAAGCTGTGGAAGTTCTTCATGGGATGACCCAGGAGGCTTCAAAGTTACTCACACAAAAGCTCCTTGGTTAAAGCCAACAACTGCTG
CAGGTGTTCATGGGATGCATTTATCTAGTCAGCAGATGTCGGACAATATTATGCCGCATGGTGCACATCATGGTTTCGATGGGTTCATGGCATTCAA
GGGAACTTGTATAAAGTTCCCTAATCAAGGTGTCCAAGCTTCTGCTGTTGCTTCCGACTCCAGTGGAGCCCCGGATCTTCAGCACGCCCTATCCCTA
CTATCCAGCAACCCAGTGGGTGCTGCCAACCTCCAGCCAAGTCCCCAGATGCACTCTGGGGTGGCAGCCATTGCCGGCGCCCCCAACCCCGCGATGC
ACGTGCTGGGATCATCGACGGGGCTCTGGCTAGACGGCGGCCAGCCCCTCGACGATCACCCGCGGTTCCAGGTCTTCGAGCGCTTGGGGGACCATGA
CAGCGAGCTCCAGCTCCCAAAGCCTTCCTACGACCATGCCTCGCACTTCGACCGGATGCAC
```

>*TaSPL4* (with disrupted miR156 binding site)

```
ATGGAGTGGACGGCCCCGAAGTCCACCCCGCTCTCCCCGCCCCACCTCCTCTGGGACTGGGGCGACGCCGCCGCGCCGGGCTCCTCCGGCGAGGCGG
CGGGGAGGCGCGGGAAGGAGAAGCGGGCCAGGGGGGAGGCGGCCGGCGGAGGAGGCGGAGGCGGAGGAGGAGGGGGCGTGAGGTGCCAGGTGGAGGG
CTGCGGGGTCGAGCTCCGCGACGCCAAGGAGTACCACCGGAAGCACCGGGTCTGCGAGGCCCACACCAAGTTCCCCCGCGTCGTCGTCGCCGGCCAG
GAGCGCCGCTTCTGCCAGCAGTGCAGCCGGTTCCATGCGCTGTCCGAGTTCGACCAGAAGAAGAGGAGCTGCCGGAGGCGGCTGTCCGATCACAATG
CTCGGCGCCGGAAACCACAGCCAGACGCATTCTCCTTTACACCGGCAAGGCTCCCGTCGTCATTGGTGTTTGATGATAGACGGCAAATAAGTTTTGT
CTGGAATAAAGATCCACTCAGCCATGCAAGGCCTTTCCCATGTTCTCCATGGGACAGCCCATCTGACTTCAAGCTCCCGCAAGTGAAGGAAATAAGA
GAAGTATCAATCAATGGACAAGTTCATTTTGATAAATCTCATCTACCCAATGCTGTTCCAGCACTGAGTCATGACATAGCTGAGCTGCTACCAATGA
AAGGTCCGGACGCATCTGTAACCGCTTCAAAATTAGGTGGAGCACCGGATCTTCAGCGCGCCCTATCCCTACTATCCGCTAGTTCTTGTGGATTACC
TGATCCTGTACAGCAAGCATCTTGTCTCGTCCAATTCACTGGTGCCAGCCAAAACAGCCGGGGCCCTTCACCACATGGAGGGAGCCCTCCATCGGCG
TCCTGTGCCGAAGGACAGCCCATGGCACCATCGCCTCAGTTCGTCCGTTTCACCATGGATGGCGCCAGCAGTGGCTATGAATCCACTTTCTTTGGCG
TAAACCGGATGAAT
```

*Electrophoretic Mobility Shift Assay (EMSA):* DNA-protein binding interactions were analyzed using the chemiluminescent EMSA kit (GS009). Biotin-labeled DNA probes and identical unlabeled (cold) competitor probes were synthesized based on two promoter regions of the *FUL-A2* promoter. The first probe (*FUL-A2*-pr.$^{-399\text{ to }-374}$) included a canonical 'GTACG' SPL-binding site, whereas the second DNA probe included a separate promoter segment (*FUL-A2*-pr.$^{-579\text{ to }-551}$) without any *SPL* binding motif. The second probe was included as a negative control. The control probes included random DNA sequences and were used as additional negative controls. All probe sequences are shown in Data S2.

The binding reaction was performed by incubating the purified protein with the binding reaction buffer at room temperature for 10 minutes. The probes were then added, and the reaction mixture was further incubated for 20 minutes at room temperature. The protein-DNA complexes were resolved on a pre-run 4% non-denaturing polyacrylamide gel in 0.5× TBE buffer at 100 V under cold conditions. The resolved complexes were subsequently transferred onto a positively charged nylon membrane using a wet transfer system at 300 mA for 45 minutes in 0.5× TBE buffer. After transfer, the DNA was cross-linked to the membrane via UV irradiation. For signal detection, the membrane was blocked, incubated with a Streptavidin-Horseradish Peroxidase (HRP) conjugate, and thoroughly washed. The biotin-labeled probes were detected using the BeyoECL Moon chemiluminescent substrate, and signals were captured by imaging.

***Method S7. Yeast-two-hybrid (Y2H) assays***

The GAL4-based Y2H system was used to investigate protein interactions. The coding sequence of *SPL3*, *SPL4* and *SPL13* without stop codon and the C-terminal GRAS domain of RHT-B1 (encoding amino acids 201-621) were cloned into the pDONRTM/Zeo vector. We then cloned these genes into Y2H vectors pGADT7 (activation-domain vector) and pGBKT7 (DNA-binding domain vector) by In-Fusion HD Cloning method. The resulting constructs were transformed into yeast AH109 gold strain using the lithium acetate method. Transformants were selected on SD medium lacking leucine (L) and tryptophan (W, abbreviated as SD-L-W). Positive transformants were replated on SD medium lacking L, W, histidine (H) and adenine (A, abbreviated as SD-L-W-H-A) to test protein-protein interactions. To test autoactivation, we co-transformed each bait vector with pGADT7 and each prey vector with pGBKT7 as a negative control.

To identify the region of SPL-A3 that interacts with DELLA, we selected two truncation mutants for our Kronos sequenced mutant population. Kronos mutant K2513 has a premature stop codon at position 282 (Q282*) that results in the truncation of 194 amino acids in the C-terminal region. This truncated peptide is designated as SPL-A3-Δ194 and is similar to the shorter SPL13 protein. Kronos mutant K1071 has a premature stop codon at position 317 (W317*) that results in the truncation of 157 amino acids in the C-terminal region. This truncated protein, designated as SPL-B3-Δ157, has 37 additional amino acids compared to SPL-A3-Δ194.

***Method S8. Bimolecular fluorescence complementation (BiFC)***

To confirm the interaction between SPL3, SPL4 and the DELLA-GRAS domain, we performed bi-molecular fluorescent complementation (BiFC or split YFP) assays. We cloned the coding sequence of *SPL3* and *SPL4* without stop codon into the 35S::NYFP-GW vector and the C-terminal GRAS domain encoded by *RHT-B1* into the 35S::CYFP-GW vector. Rice protoplasts were prepared from the leaves of the variety Kitaake, transfected and visualized as described in a previously published study (He *et al.*, 2016). We co-transformed 35S::CYFP- RHT1-GRAS with either 35S::NYFP-SPL3 or 35S::NYFP-SPL4 into rice protoplasts and documented the presence or absence of fluorescent nuclear signals. The CYFP empty vector was used as a negative control for 35S::NYFP-SPL3 and -SPL4 constructs and the NYFP empty vector was used as a negative control for 35S::CYFP- RHT1-GRAS.

***Method S9. CoIP assays***

The coding sequences (CDs) of *VRN1*, *FUL2*, *SPL-B3* and *SPL-A4* were synthesized and inserted into the Super1300-FLAG vector (Chen *et al.*, 2023). Seven silent mutations were introduced in the miR156 binding site of SPL3 and SPL4 by overlapping PCR to generate miR156 resistant alleles (Primers in Data S2). These mutations are important because protoplasts are prepared from seedlings, where miR156 expression is high. The modified bases are indicated in red letters and yellow highlight in the sequences below. Co-transformations of DELLA-

GRAS-GFP with either SPL3-FLAG or SPL4-FLAG were done in *N. benthamiana* leaves, and after immunoprecipitation with anti-GFP beads, SPL3-FLAG and SPL4-FLAG were detected using an anti-FLAG antibody.

The CDs of DELLA- GRAS (encoding amino acids 201-621) were synthesized (sequence below) and inserted into the Super1300-GFP vector (Jiang *et al.*, 2023). These constructs were transformed into *Agrobacterium tumefaciens* GV3101 and used for transient expression in *N. benthamiana*. The DELLA-GRAS-GFP was co-transformed with either VRN1-FLAG or FUL2-FLAG into *N. benthamiana* leaves, and after immunoprecipitation with anti-GFP beads, the presence of VRN1-FLAG and FUL2-FLAG was tested with an anti-FLAG antibody.

Total proteins were extracted at 48 h post infiltration and then incubated with GFP-tagged magnetic beads. The beads were collected by centrifugation and then washed five times with cold 1x TBS buffer (containing 0.5% Triton X-100). Proteins were released from the beads by incubating at 100 ˚C for 8 min with 50 µL 1x TBS. Immune precipitates were separated by SDS–PAGE gels and detected by immunoblotting with monoclonal α-GFP and α-Flag antibodies (Abmart Inc.). The coIP experiments were conducted by ProNet Biotech Co., Ltd. (https://www.pronetbio.com/, Jiangsu, China).

**Sequences**

Silent mutations introduced in the miR156 binding site to generate miR156 resistant alleles are indicated in red and highlighted in yellow.

>r*SPL-B3* Kronos coding sequence (JD06)
ATGGGCTCTTTTGGGATGGAGTGGAACCAGAGGAGCTCGGTGCTGTGGGACTGGGAGAATTTCCCGCCGATAGGCAATAACCCGAAGAACGCGATGC
AGGCTGATCCAAGATTCGCCGGCGTTGCGGCTACCATGGGGAATGAACCGCTCCATTCTTCTGGCGGCAGCGGCACCTTCTCTTCCAGCTCGGAGAT
GGGGTATGGCTCTTCCAAGAGCTCCATGTCCGCGTCGATCGATTCTTCGTCCAGGGCTGGCAACAACATGGAGTTCAGATTTGCGCCTGTCAAAAAC
CCTGACAGGAACACGAGCAAAAACCCTGAGTTGGGTAAAGTTGACAACACGAGGACTGGAACATCTCCGTCGTCTGTGGTGGCAGTGAGCAGTGGAG
AGCCGGTGATCGGCCTGAAGCTTGGCAAGAGAACTTACTTCGAGGATGTCTGTGGAGGGCAGAGTGTCAAGAGCTTGCCATCGGGTGCTGCGAGCGC
GCCAAACAAATCTCCTGCTTTGGGCAAGAAGGCAAAGGCGGAACAACAGAAGCCACATAACTCATACTGTCAGGTTGAAGGCTGCAAAGCCGATCTC
TCTTCTGTTAAAGATTACCATAGGAAGCACAGGGTCTGTGAACTTCACTCTAAGGCTCCCAAAGTTGTTGTCGCTGGTCTGGAGCGACGCTTTTGCC
AACAGTGCAGCCGGTTTCATGCTTTAGCTGAGTTTGACCAGAAAAAGCGAAGCTGCCGTAGGCGTCTCAATGATCATAATTCCCGCAGGCGCAAGCC
ACAGCCAGAAGCAATTTCTTTCAGTTCATCAAGGATGTCTACGATGTTTTATGATGCAAGGCAACAGCCTAATTTCCTATTTGGTCAGGCTCCTTAT
GTTCAAATGAGAAGCTGTGGAAGTTCTTCATGGGATGACCCAGGAGGCTTCAAAGTTACTCACACAAAAGCTCCTTGGTTAAAGCCAACAACTGCTG
CAGGTGTTCATGGGATGCATTTATCTAGTCAGCAGATGTCGGACAATATTATGCCGCATGGTGCACATCATGGTTTCGATGGGTTCATGGCATTCAA
GGGAACTTGTATAAAGTTCCCTAATCAAGGTGTCCAAGCTTCTGCTGTTGCTTCCGACTCCAGTGGAGCCCCGGATCTTCAGCACGCCCTATCCCTA
CTATCCAGCAACCCAGTGGGTGCTGCCAACCTCCAGCCAAGTCCCCAGATGCACTCTGGGGTGGCAGCCATTGCCGGCGCCCCCAACCCCGCGATGC
ACGTGCTGGGATCATCGACGGGGCTCTGGCTAGACGGCGGCCAGCCCCTCGACGATCACCCGCGGTTCCAGGTCTTCGAGCGCTTGGGGGACCATGA
CAGCGAGCTCCAGCTCCCAAAGCCTTCCTACGACCATGCCTCGCACTTCGACCGGATGCAC

>*rSPL-A4* coding sequence (JD362)
ATGGAGTGGACGGCCCCGAAGTCCACCCCGCTCTCCCCGCCCCACCTCCTCTGGGACTGGGGCGACGCCGCCGCGCCGGGCTCCTCCGGCGAGGCGG
CGGGGAGGCGCGGGAAGGAGAAGCGGGCCAGGGGGGAGGCGGCCGGCGGAGGAGGCGGAGGCGGAGGAGGAGGGGGCGTGAGGTGCCAGGTGGAGGG
CTGCGGGGTCGAGCTCCGCGACGCCAAGGAGTACCACCGGAAGCACCGGGTCTGCGAGGCCCACACCAAGTTCCCCCGCGTCGTCGTCGCCGGCCAG
GAGCGCCGCTTCTGCCAGCAGTGCAGCCGGTTCCATGCGCTGTCCGAGTTCGACCAGAAGAAGAGGAGCTGCCGGAGGCGGCTGTCCGATCACAATG
CTCGGCGCCGGAAACCACAGCCAGACGCATTCTCCTTTACACCGGCAAGGCTCCCGTCGTCATTGGTGTTTGATGATAGACGGCAAATAAGTTTTGT
CTGGAATAAAGATCCACTCAGCCATGCAAGGCCTTTCCCATGTTCTCCATGGGACAGCCCATCTGACTTCAAGCTCCCGCAAGTGAAGGAAATAAGA
GAAGTATCAATCAATGGACAAGTTCATTTTGATAAATCTCATCTACCCAATGCTGTTCCAGCACTGAGTCATGACATAGCTGAGCTGCTACCAATGA
AAGGTCCGGACGCATCTGTAACCGCTTCAAAATTAGGTGGAGCACCGGATCTTCAGCGCGCCCTATCCCTACTATCCGCTAGTTCTTGTGGATTACC
TGATCCTGTACAGCAAGCATCTTGTCTCGTCCAATTCACTGGTGCCAGCCAAAACAGCCGGGGCCCTTCACCACATGGAGGGAGCCCTCCATCGGCG

```
TCCTGTGCCGAAGGACAGCCCATGGCACCATCGCCTCAGTTCGTCCGTTTCACCATGGATGGCGCCAGCAGTGGCTATGAATCCACTTTCTTTGGCG
TAAACCGGATGAAT

>DELLA-GRASS coding sequence TraesCS4B02G043100.1 Rht-B1
ATGAGCTCTGTGGTGGAGGCTGCCCCGCCGGTGGCCGCCGCGGCCGGTGCGCCCGCGCTGCCGGTCGTCGTGGTCGACACGCAGGAGGCCGGGATTC
GGCTGGTGCACGCGCTGCTGGCGTGCGCAGAGGCCGTGCAGCAGGAGAACTTCTCTGCCGCGGAGGCGCTGGTGAAGCAGATACCCTTGCTGGCCGC
GTCCCAGGGCGGCGCCATGCGCAAGGTCGCCGCCTACTTCGGCGAGGCCCTCGCCCGCCGCGTCTTCCGCTTCCGCCCGCAGCCGGACAGCTCCCTC
CTCGACGCCGCCTTCGCCGACCTCCTCCACGCGCACTTCTACGAGTCCTGCCCCTACCTCAAGTTCGCCCACTTCACCGCCAACCAGGCCATCCTGG
AGGCGTTCGCCGGCTGCCGCCGCGTGCACGTCGTCGACTTCGGCATCAAGCAGGGGATGCAGTGGCCCGCCCTTCTCCAGGCCCTGGCGCTCCGTCC
CGGCGGCCCTCCCTCGTTCCGCCTCACCGGCGTCGGCCCCCCGCAGCCGGACGAGACCGACGCCTTGCAGCAGGTGGGCTGGAAGCTCGCCCAGTTC
GCGCACACCATCCGCGTCGACTTCCAGTACCGCGGCCTCGTCGCCGCCACGCTCGCGGACCTGGAGCCGTTCATGCTGCAGCCGGAGGGCGAGGAGG
ACCCGAACGAGGAGCCCGAGGTAATCGCCGTCAACTCGGTCTTCGAGATGCACCGGCTGCTCGCGCAGCCCGGCGCCCTGGAGAAGGTCCTGGGCAC
CGTGCGCGCCGTGCGGCCGAGGATCGTCACCGTGGTGGAGCAGGAGGCGAACCACAACTCCGGCACATTCCTGGACCGCTTCACCGAGTCCCTGCAC
TACTACTCCACCATGTTCGATTCTCTGGAGGGCGGCAGCTCCGGCGGCCCATCCGAAGTCTCATCTGGGGCGGCTGCTGCTCCTGCCGCCGCCGGCA
CGGACCAGGTCATGTCCGAGGTGTACCTCGGCCGGCAGATCTGCAACGTGGTGGCCTGCGAGGGGGCGGAGCGCACAGAGCGGCACGAGACCCTGGG
GCAGTGGCGGAACCGCCTCGGCAACGCCGGGTTCGAGACCGTCCACCTGGGCTCCAATGCCTACAAGCAGGCGAGCACGCTGCTGGCGCTCTTCGCA
GGCGGCGACGGGTACAAGGTGGAGGAGAAGGAGGGCTGCCTGACGCTGGGGTGGCACACGCGCCCGCTGATCGCCACCTCCGCATGGCGCCTGGCCG
CGCCG
```

#### *Method S10. Yeast three-hybrid (Y3H) assays*

The pBridge yeast three-hybrid system (Clontech) was employed for the Y3H assays. This assay can test the effect of a bridge protein, referred to as the 3$^{rd}$ protein in this study, on the interaction between two proteins, one fused to a DNA-binding domain (BD) and the other to an activation domain (AD). This 3$^{rd}$ protein is driven by an inducible MET25 promoter and can only be expressed in the absence of methionine. The addition of 1 mM methionine to the medium is sufficient to inhibit its expression. We generated four Y3H bait constructs, including pBridge-SPL3-pMET25-VRN1, pBridge-SPL3-pMET25-FUL2, pBridge-SPL4-pMET25-VRN1, and pBridge-SPL4-pMET25-FUL2 using a two-step cloning strategy and primers described in Data S2.

In the first step, the coding region of *SPL3* or *SPL4* was inserted downstream of the GAL4-BD in the pBridge vector between the *Eco*RI and *Bam*HI restriction sites. In the second step, the coding region of *VRN1* or *FUL2* was cloned downstream of the MET25 promoter between the *Not*I and *Bgl*II restriction sites. The C-terminal GRAS domain of the *RHT1* gene (encoding amino acids 201-621) was cloned into the pGADT7 vector between the *Nde*I and *Eco*RI restriction sites to generate the prey construct pGADT7-RHT1-GRAS. Each of the four pBridge bait constructs was then paired with the pGADT7-RHT1-GRAS prey vector and co-transformed into Y2H Gold yeast strain (Clontech). Transformants containing both vectors were selected on SD medium lacking leucine (L) and tryptophan (W) medium (SD-L-W). Protein interactions were quantified using quantitative alpha-galactosidase assays, as previously described (Li *et al.*, 2011).

#### *Method S11. Statistical analyses*

Statistical analyses were performed using SAS version 9.4. The ANOVA assumption of homogeneity of variances was tested using the Levene's test and the assumption of normality of residuals using the Shapiro-Wilk test. When these assumptions were not met, data were transformed using different power transformations until the assumptions were satisfied. The transformations used are indicated in the respective analyses. When none of the transformations was effective in restoring the ANOVA assumptions, we used the Kruskal-Wallis non-parametric test. Graphs and tables present the untransformed data, whereas the *P* values correspond to the transformed data. The significance of the genetic interactions was tested using factorial ANOVAs, with genes treated as factors and alleles as different levels within each factor.

We used boxplots in most of the graphs to show both the means and distribution of the data. In these graphs, the box shows the range from first to third quartiles and is divided by the median. The whiskers span down to the minimum, and up to the maximum observation. Results from individual experiments are indicated by empty black circles.
