## Supplemental Figures for "MicroRNA156 and its targeted *SPL* genes interact with the photoperiod, vernalization, and gibberellin pathways to regulate wheat heading time"

### Supplementary Figures

**Figure S1**. miR156 binding sites in nine wheat *SPL* genes and their expression profiles.

(**a**) miR156 binding sites of wheat *SPL* genes. Green letters represent natural polymorphisms within the conserved miR156 binding site.

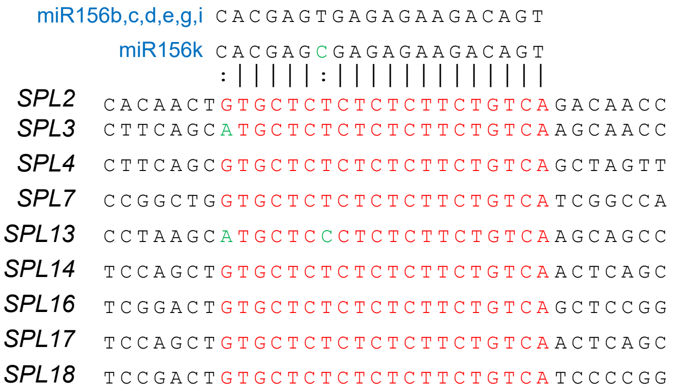


(**b**) Expression profiles of the A, B and D homeologs of the nine wheat *SPL* genes with miR156 binding sites based on ExpVIP (http://www.wheat-expression.com/, $\log_2$ TPM). The three rows in root and leaf/shoot represent samples collected at seedling, vegetative and reproductive stages, respectively. This graph is based on four RNA-seq studies in hexaploid wheat selected from ExpVIP: Developmental time courses in CS and Azhurnaya, CS leaves and roots from 7 leaf stages and CS seedlings (leaf and roots) and developing spikes.

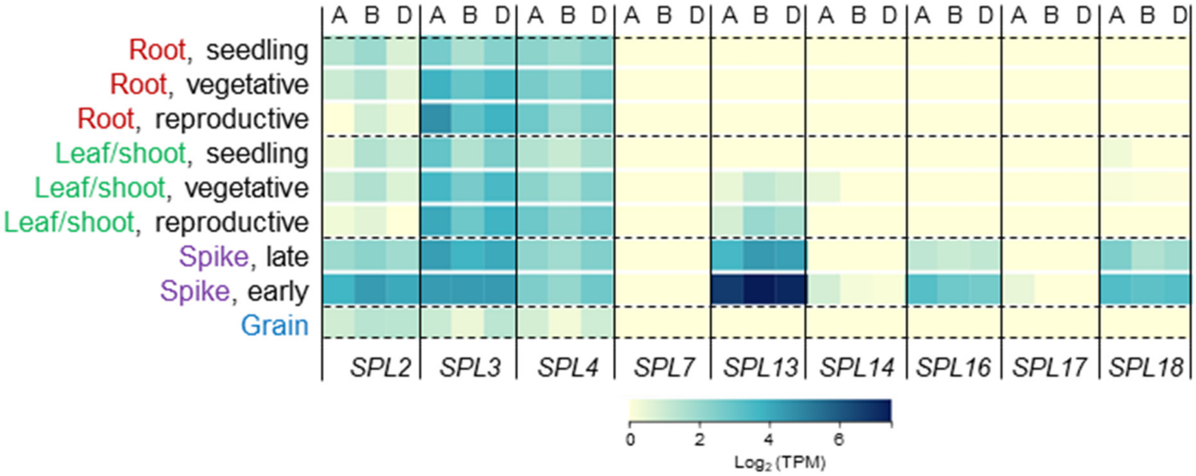

**Figure S1**. Continuation.

(**c**) Single molecule fluorescent in situ hybridization (smFISH) of miR156 regulated SPLs at the transition of the shoot apical meristem (SAM) into an inflorescence meristem (IM) in tetraploid wheat Kronos. Data was obtained from X. Xu et al. (2025). Genome Biology 26:352. [https://doi.org/10.1186/s13059-025-03811-3](https://doi.org/10.1186/s13059-025-03811-3)

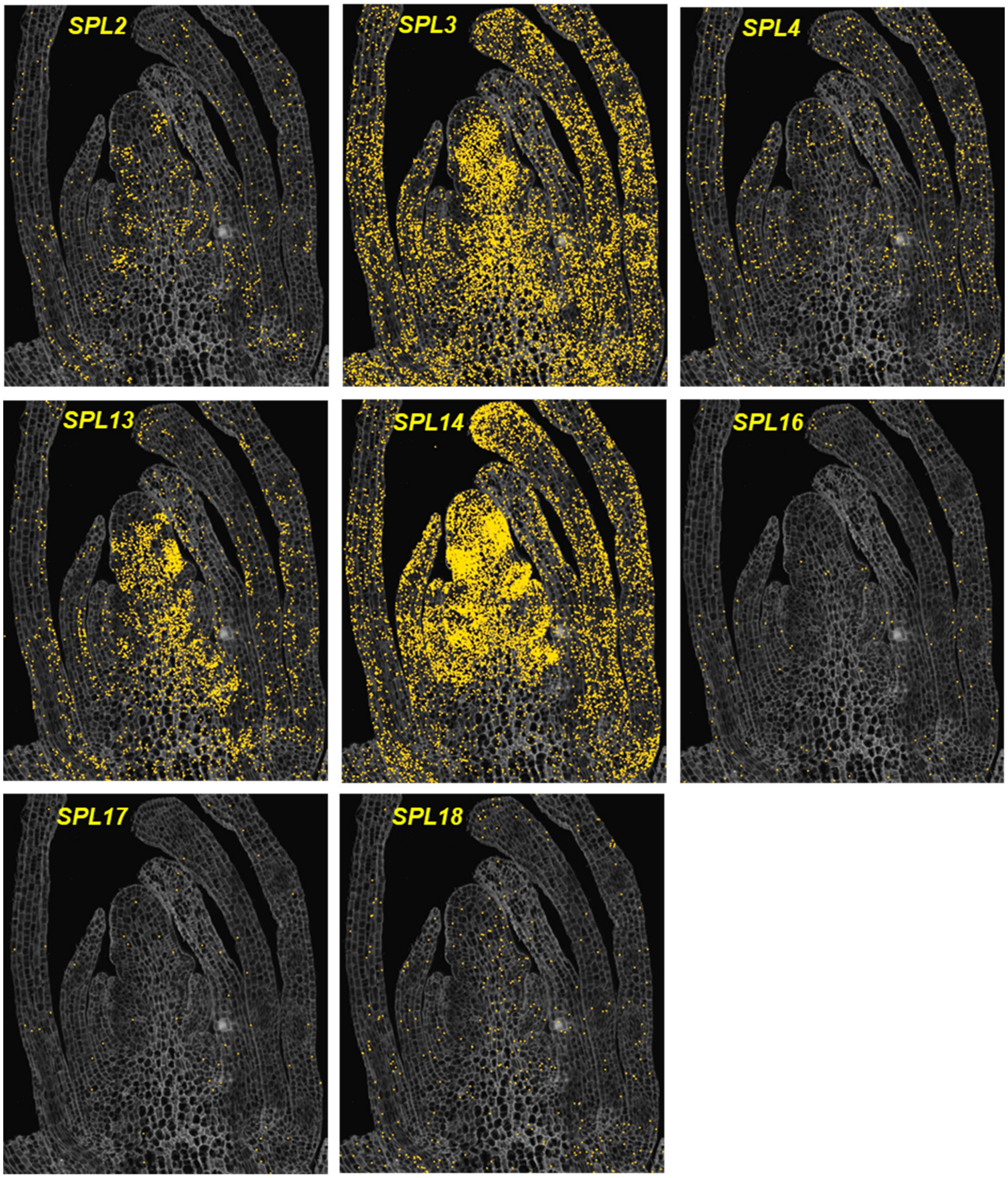

**Figure S2.** Alignment of SBP domains for Arabidopsis, rice and wheat SPL proteins.

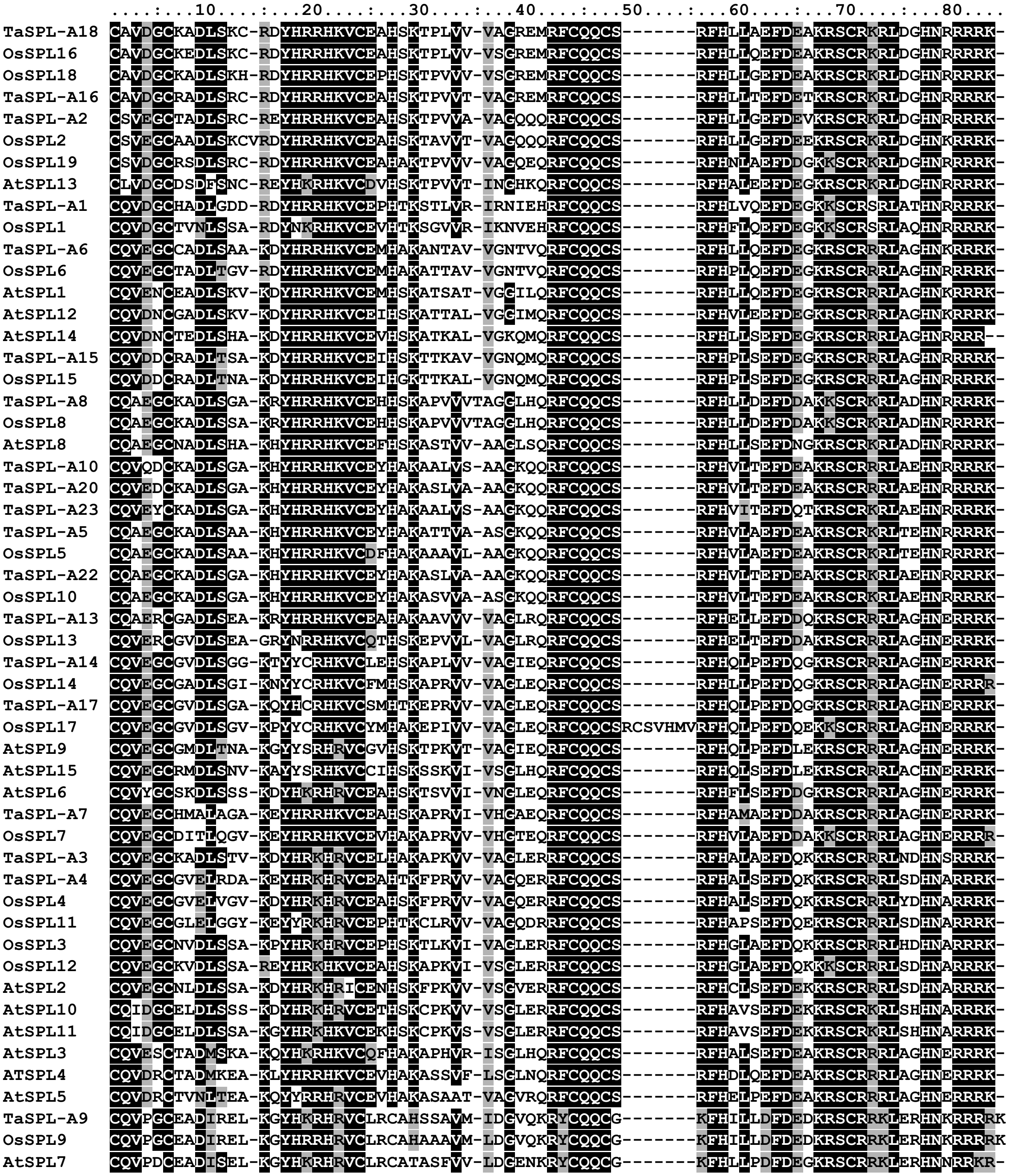

**Figure S3.** Phylogenetic tree of SPL proteins.

Phylogenetic tree based on SBP alignment (Figure S2, Method S1) using the Maximum Likelihood method and Le_Gascuel_2008 model (Le and Gascuel, 2008).

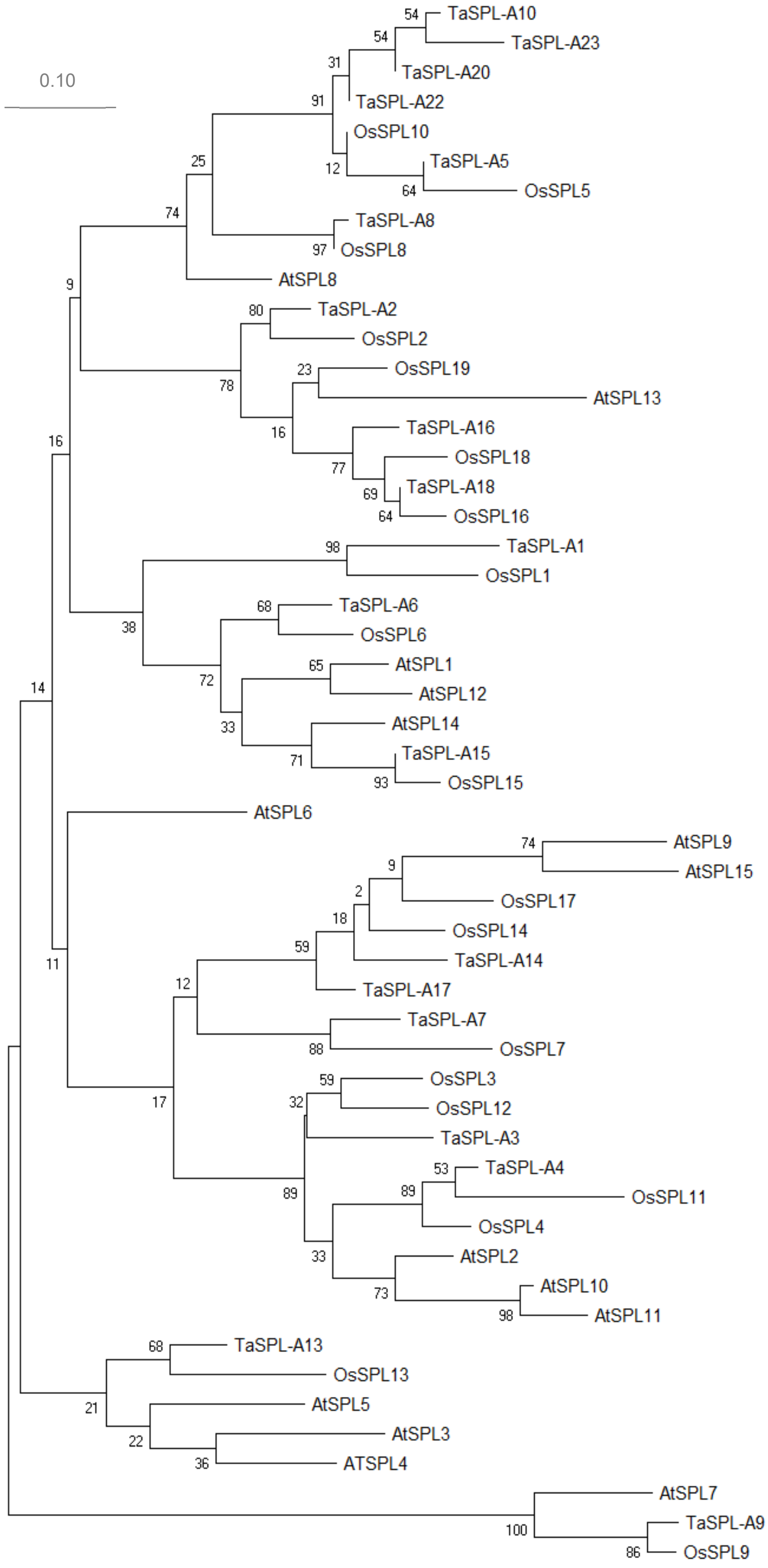


The percentage of trees in which the associated taxa clustered together is shown next to the branches. Initial tree(s) for the heuristic search were obtained the Neighbor-Join and BioNJ algorithms to a matrix of pairwise distances and then selecting the topology with superior log likelihood value. The tree is drawn to scale, with branch lengths measured in the number of substitutions per site. This analysis involved 53 amino acid sequences in the SBP domain (Figure S2). Evolutionary analyses were conducted in MEGA X (Kumar *et al.*, 2009).

**Figure S4**. Regulation of *SPL3*, *SPL4*, *SPL13* and *SPL2* expression by miR156.

**(a-d)** Boxplots showing expression of *SPL3*, *SPL4*, *SPL13* and *SPL2* in leaves 1, 5 and 9 (n= 4). **(e-h)** Effect of overexpression of miR156 on *SPL3*, *SPL4*, *SPL13* and *SPL2* expression (n= 6). Overexpression of miR156 occurs only when the pOp::miR156 and the Ubi::LhG4 are combined by crossing. **(i-l)** Effect of MIM156 on the transcript levels of *SPL3*, *SPL4*, *SPL13* and *SPL2* in five independent transgenic events. **(a-h)** Different letters indicate significant differences in Tukey tests ($P < 0.05$). Transcript levels are expressed relative to *ACTIN* and were calculated using the ΔCt method. **(i-l)** Dunnett tests between transgenics and wildtype, n= 4. ns= not significant, * = $P < 0.05$, ** = $P < 0.01$, *** = $P < 0.001$. Raw data and statistical analyses are available in Data S3.

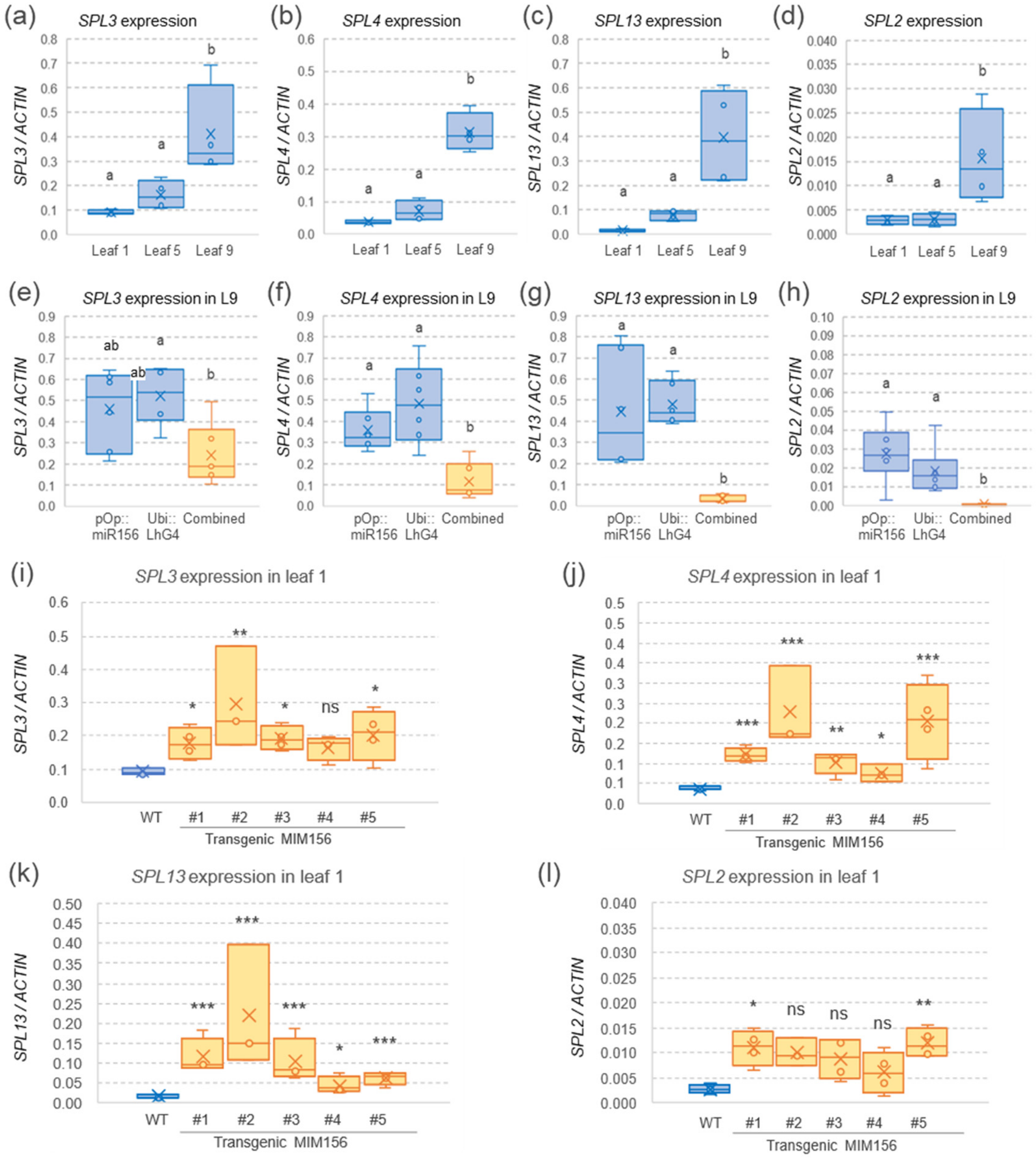

**Figure S5**. Expression of wildtype and resistant alleles *rSPL3*, *rSPL4*, and *rSPL13*.

**(a)** *rSPL-A3*, **(b)** *rSPL-B3*, **(c)** *rSPL-A4*, **(d)** *rSPL-A13* and **(e)** *rSPL-B13.* Transcript levels are expressed relative to *ACTIN* and were calculated using the ΔCt method. Probability values correspond to *t*-tests between resistant and wildtype alleles. ns= not significant, *= *P*< 0.05, **= *P*< 0.01, ***= *P*< 0.001. n= 6 plants per genotype (except for *rSPL-A4*, n=4). Raw data and statistical analyses are available in Data S3.

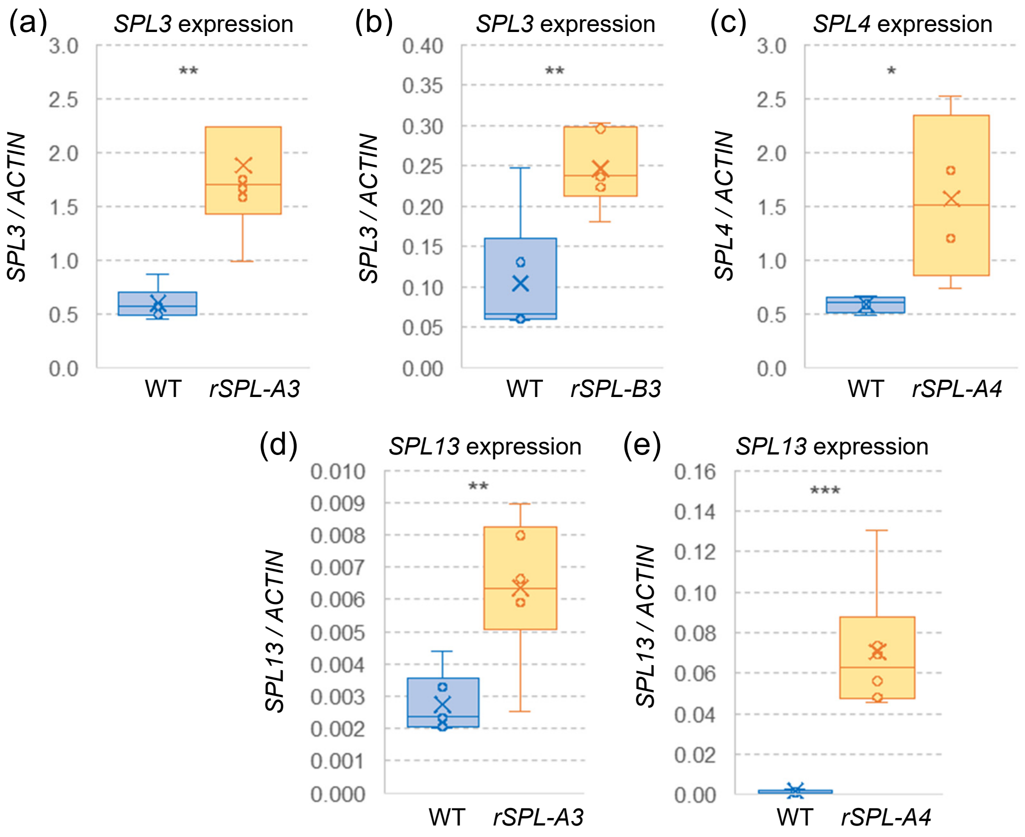

**Figure S6**. Pictures of wildtype and *rSPL3*, *rSPL4*, and *rSPL13* single mutants at heading time.

(**a**) *rSPL-A3*. (**b**) *rSPL-B3*. (**c**) *rSPL-A4*. (**d**) *rSPL-A13*. (**e**) *rSPL-B13*. Pictures were taken at the time of spike emergence in the mutant lines. Scale bar is 5 cm. Arrows at the top of the main tillers show emerging spikes in the *rSPL* mutants but not in their respective wildtype sister lines.

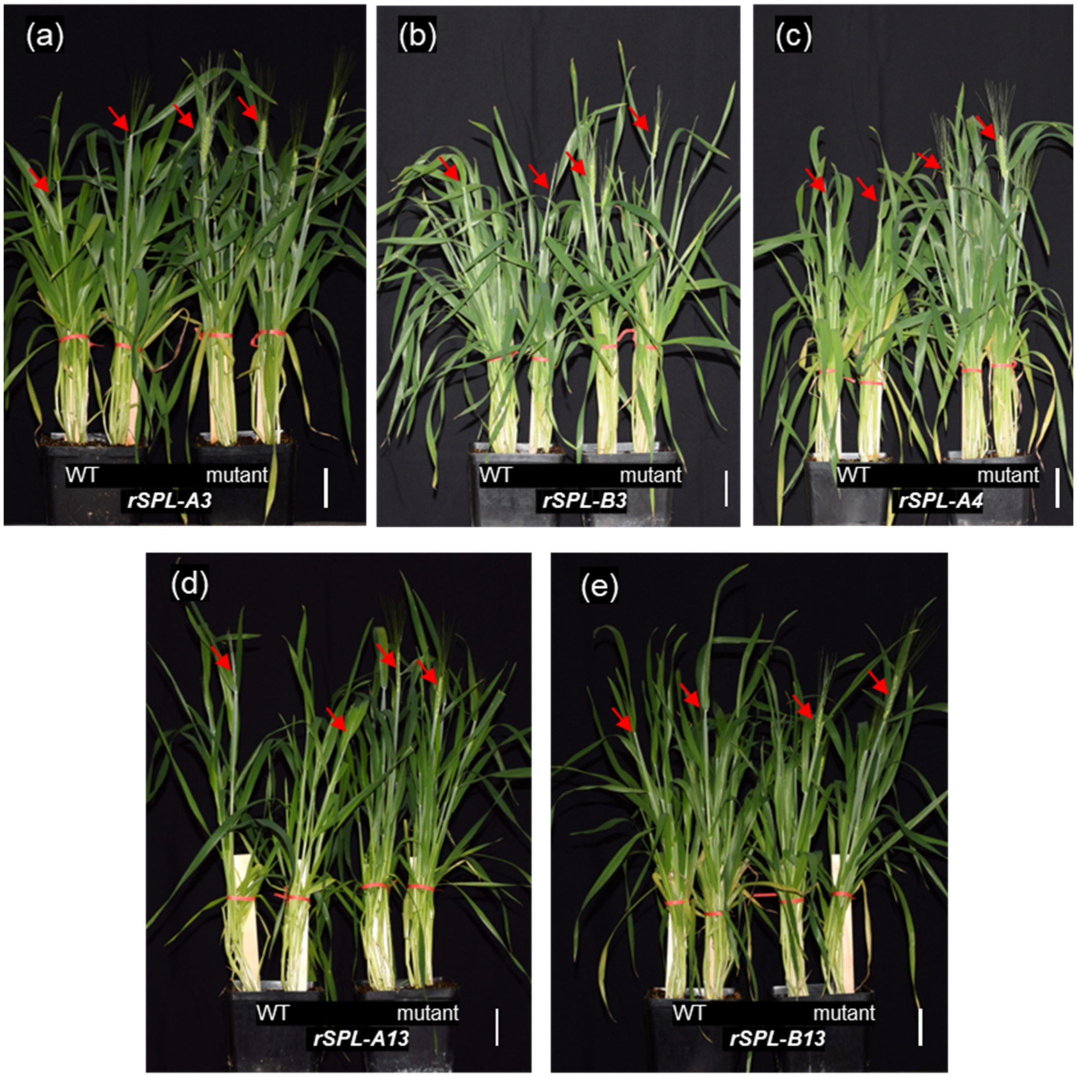

**Figure S7**. Effect of *rSPL3*, *rSPL4*, *rSPL13* on leaf number.

**(a)** Leaf number of WT (n= 10) and r*SPL-A3* (n= 5). **(b)** Leaf number of WT (n= 14) and *rSPL-B3* (n= 9). **(c)** Leaf number of WT (n= 7) and *rSPL-B13* (n= 8). **(d)** Leaf number of single *rSPL-A4* and *rSPL-A13* and combined mutants compared with wildtype (n= 12). **(e)** Leaf number of triple resistant mutant *rSPL-B3/A4/A13* (n= 15) compared with WT (n= 14). ns = not significant, *= $P < 0.05$, **= $P < 0.01$, ***= $P < 0.001$. $P$ values are from non-parametric Kruskal-Wallis test *vs.* wildtype. Non-parametric tests were used due to the lack of normality. Raw data and statistical analyses are available in Data S5.

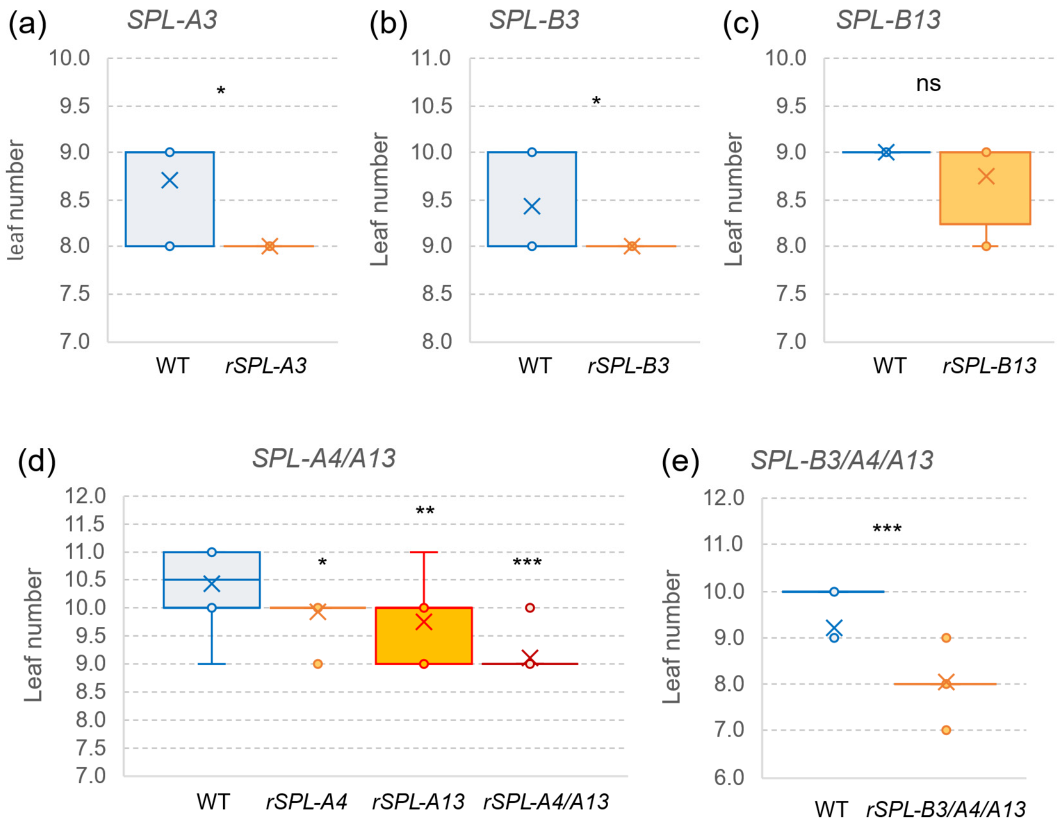

**Figure S8**. Effect of *rSP3*, *rSPL4*, and *rSPL13* alleles on heading time under LD and SD.

Interaction graphs for heading time of plants carrying different *SPL* resistant allele combinations grown under LD (16h light and 8 h darkness) and SD (8h light and 16 h darkness) conditions. **(a)** *rSPL-A4*, experiment 1. **(b)** *rSPL-A4*, experiment 2. **(c)** *rSPL-B3*. **(d)** *rSPL-A13*. **(e)** combined *rSPL-A4 rSPL-A13*. **(f)** combined *rSPL-B3 rSPL-A4 rSPL-A13*. Raw data and statistical analyses are available in Data S7.

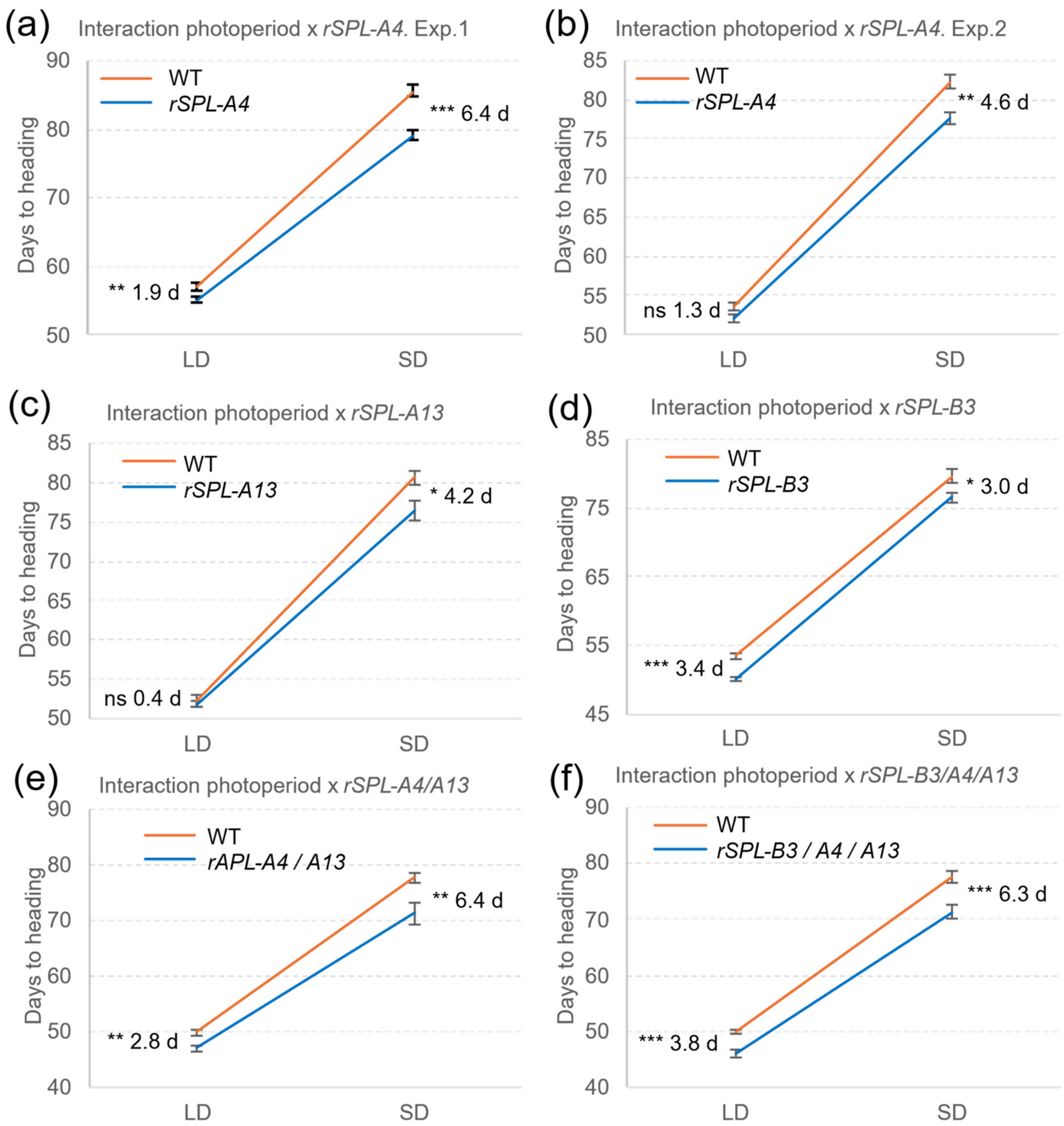

**Figure S9.** Yeast-one-hybrid assays (Y1H, Method S5).

(**a**) Bait *VRN-A1* promoter (-404 to -350). (**b**) Bait *VRN-B1* intron (+190 +244). (**c**) Bait *FUL-A2* promoter (-836 to -666). (**d**) Bait *FUL-B2* promoter (-992 -826). (**e**) Bait *FUL-A2* intron (+2,556 to +2,652). (**f**) Bait *FUL-B2* intron (+1,347 to +1,514).

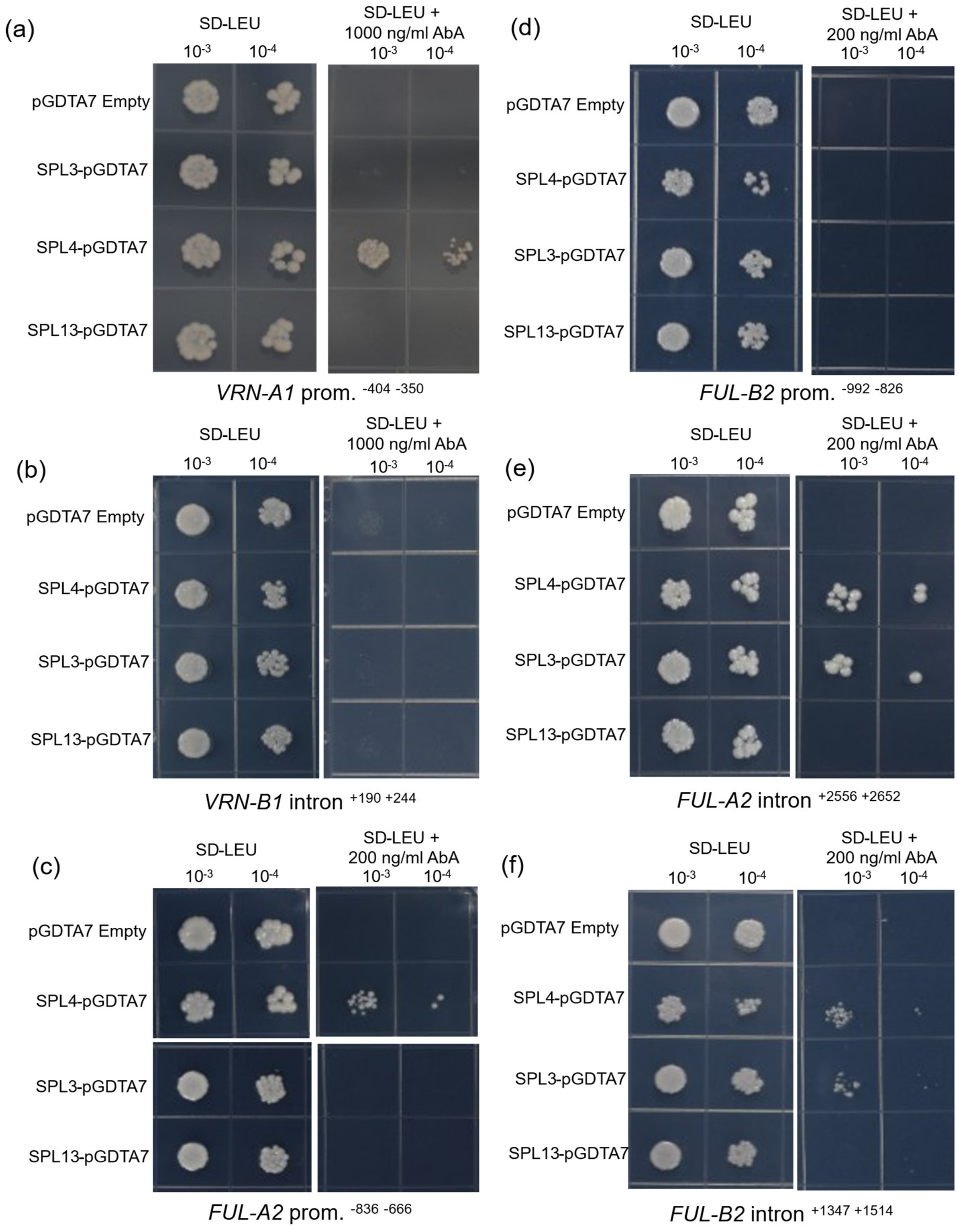

**Figure S10.** Electrophoretic Mobility Shift Assay (EMSA, Method S6).

(**a**) SDS-PAGE analysis of GST-super1300-SPL fusion protein. M: Marker; 1: GST-SPL (input); 2: GST-SPL (purified). Yellow arrowheads indicate the tagged SPL protein. (**b-d**) Binding of GST-tagged SPL proteins (GST-SPL) to a biotin-labeled DNA probe including a segment of the wheat *FUL-A2* promoter (*FUL-A2*-pr.$^{-399 \text{ to } -374}$) with a canonical 'GTACG' SPL-binding site (left panels). A second DNA segment of the *FUL-A2* promoter (*FUL-A2*-pr.$^{-579 \text{ to } -551}$) without any *SPL* binding motif was used as a negative control (right panels). Cold probes with the same DNA sequence as the probes but without the biotin label were added at 10x and 50X concentrations to test competition with the labelled probe. The control probe had a random DNA sequence (Data S2) and was included as a second negative control. Black arrows indicate the position of the shifted SPL-DNA complex, with the free probe shown at the bottom. (**b**) SPL3, (**c**) SPL4, (**d**) SPL13.

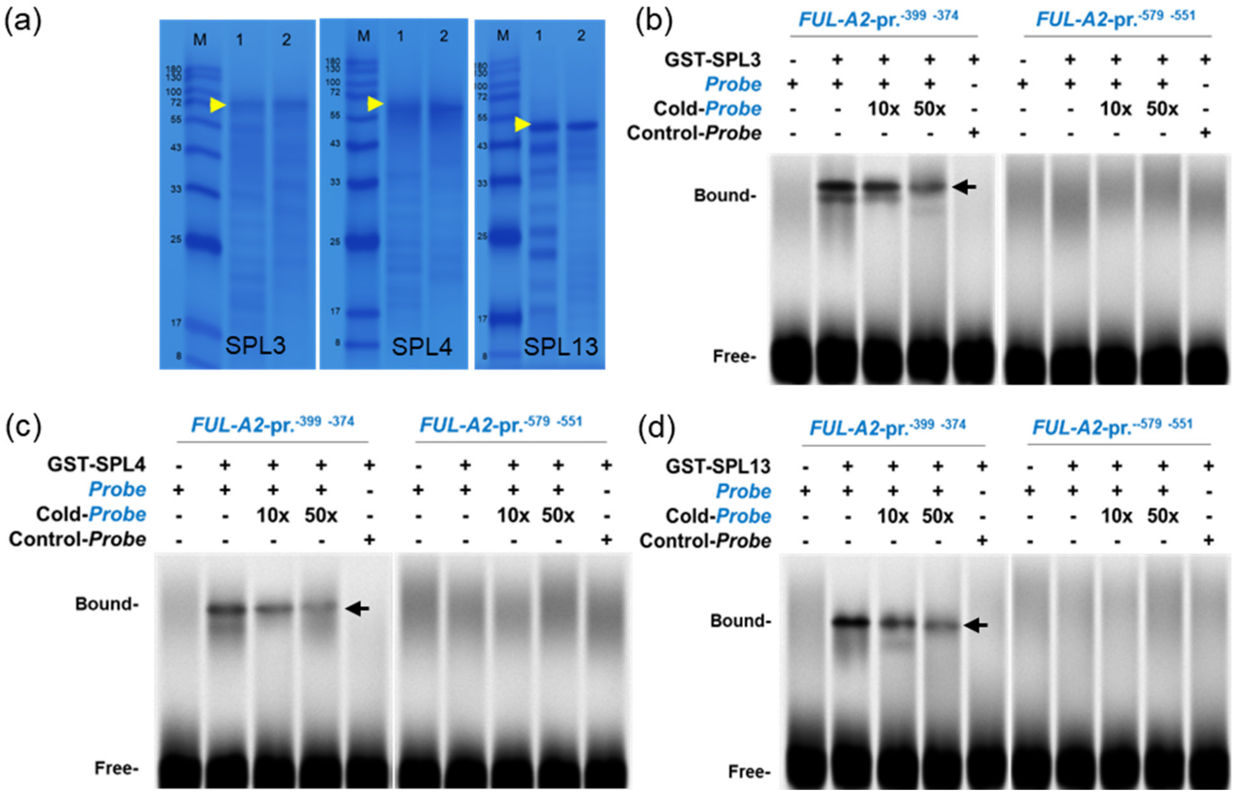

**Figure S11.** Interactions between DELLA and SPL proteins (Y2H, Method S7).

**(a)** Yeast-two-hybrid (Y2H) assays showing interactions between a truncated RHT1-GRAS and both SPL3 and SPL4 proteins**. (b)** Y2H assays showing no interactions between a truncated RHT1-GRAS and both SPL13 and a negative control with an empty vector. **(c)** Schematic representation of the three SPL proteins. Green indicates the conserved SBP domain and red the miR156 binding site. **(d)** Schematic representation of two truncated SPL3 proteins lacking 157 or 194 amino acids in the C-terminal region. **(e)** Y2H interactions between DELLA-GRAS and SPL3 truncations Δ157 and Δ194. **(f)** T-COFFEE (v11.0) alignment of partial sequences of wheat SPL3 and SPL4 with Arabidopsis AtSPL2, AtSPL10, and AtSPL11 (these proteins are known to interact with DELLA). The conserved SBP domain and the region critical for the interaction with DELLA are indicated in pink (see color scale below sequence).

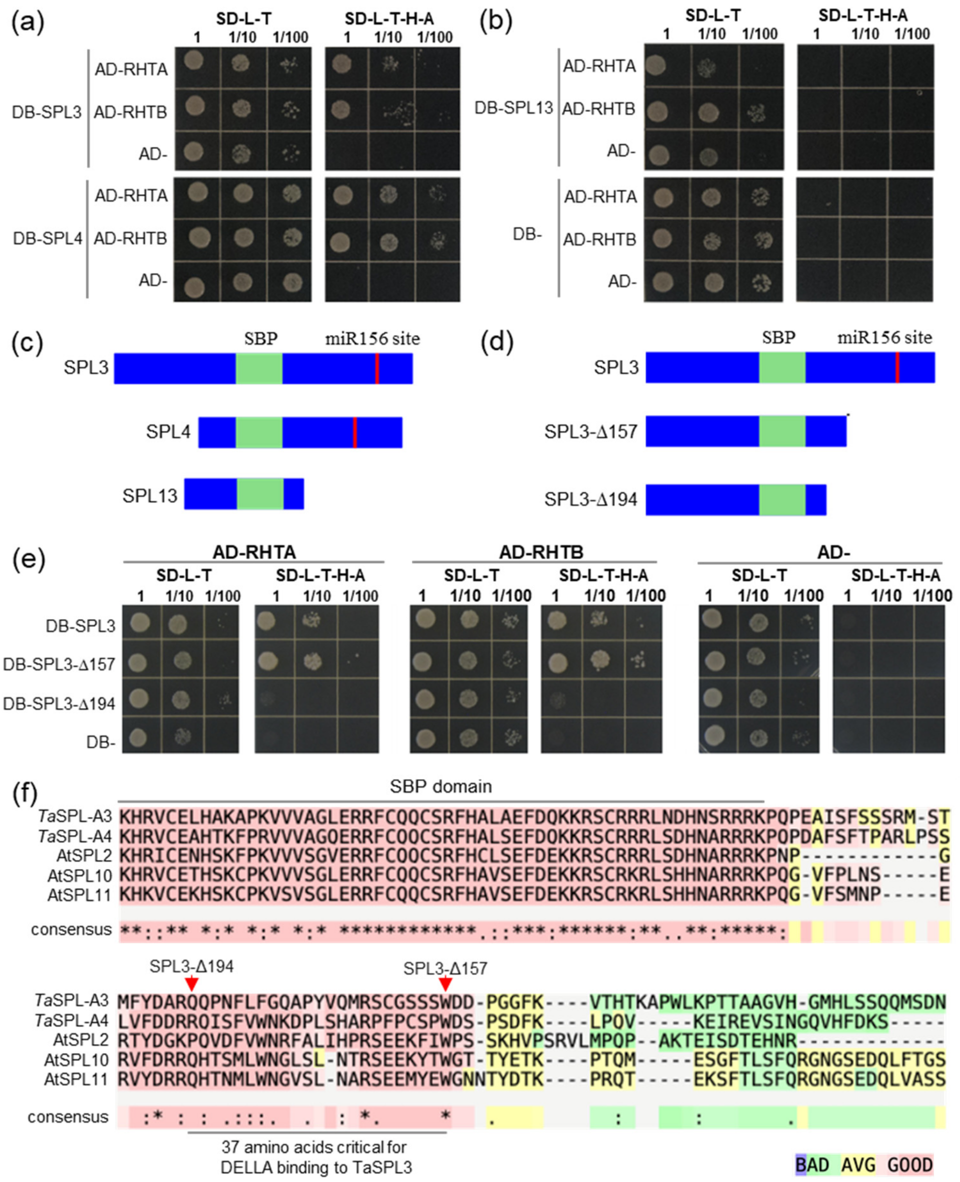

**Figure S12.** Bi-molecular fluorescent complementation (BiFC or split YFP).

BiFC assays testing the interactions between DELLA-GRASS (= RHT1) and both SPL3 and SPL4 proteins in rice protoplasts. Two independent experimental replications are presented. The three left columns correspond to the 35S::CYFP-RHT1-GRAS vector and the three right columns to the empty CYFP vector (negative control). Each set of three columns includes the YFP, brightfield, and merged pictures. The first two rows are two replications of the co-transformed empty NYFP vector (additional negative control), followed by two replications of the co-transformed 35S::NYFP-SPL3 vector and two replications of the co-transformed 35S::NYFP-SPL4 vector. Bars are 20 µm. No fluorescence signal was detected for the different combinations of empty vectors used as negative controls. See Methods S8.

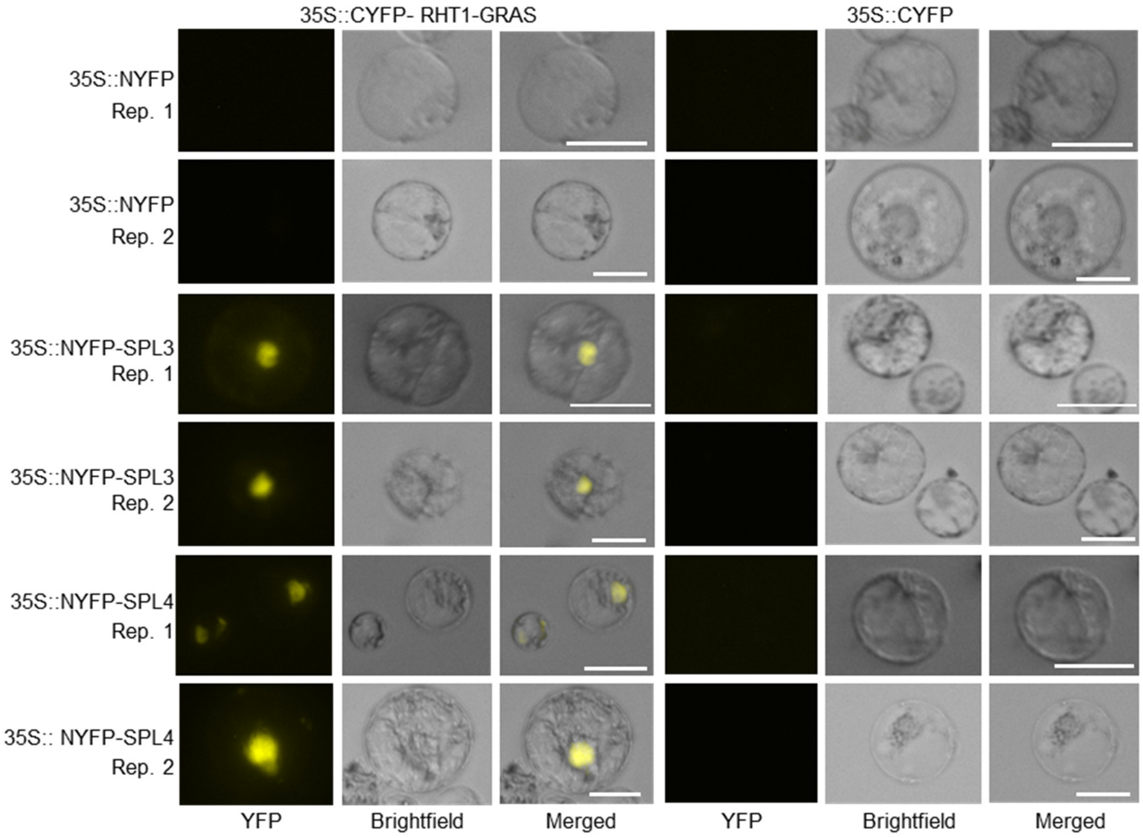

**Figure S13**. Co-IP experiments showing interactions between DELLA and SPL3 /SPL4 proteins.

Co-IP assays between DELLA-GRASS (= RHT1) and both SPL3 **(a)** and SPL4 **(b)** proteins in *N. benthamiana*. Seven silent mutations were introduced in the miR156 binding site of *SPL3* and *SPL4* to generate miR156-resistant alleles (see Method S9). Co-IP experiments were performed using GFP-tagged magnetic beads. DELLA-GRASS and the GFP empty vector were detected with anti-GFP antibody, while SPL3 and SPL4 were detected with anti-FLAG antibody. The GFP and FLAG empty vectors served as negative controls.

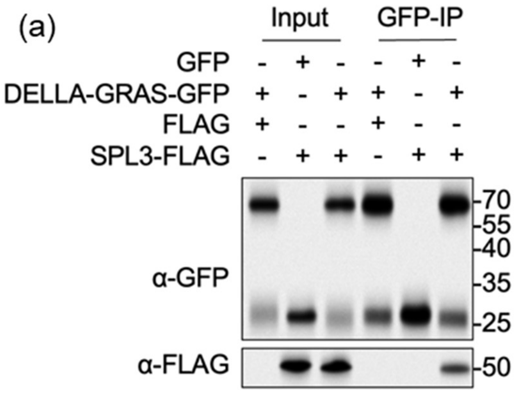

**Figure S14**. Effects of miR156 and *RHT-B1* on gene expression.

Expression analysis of **(a)** miR156, **(b)** miR172, **(c)** *SPL3*, **(d)** *SPL4*, **(e)** *SPL13* in the third leaves of WT (*Rht-A1a Rht-B1b*, GA-insensitive), MIM156, *Rht-A1a rht-B1*-null (GA-sensitive), and double mutants. Expression was determined by qRT-PCR using *SnoR* as endogenous control for miR156 and miR172 and *ACTIN* for *SPL3*, *SPL4*, and *SPL13*. *P* values are based on Dunnett tests against wildtype (WT). ns= not significant, *= $P< 0.05$, **= $P< 0.01$, ***= $P< 0.001$. Raw data and statistical analyses are available in Data S12.

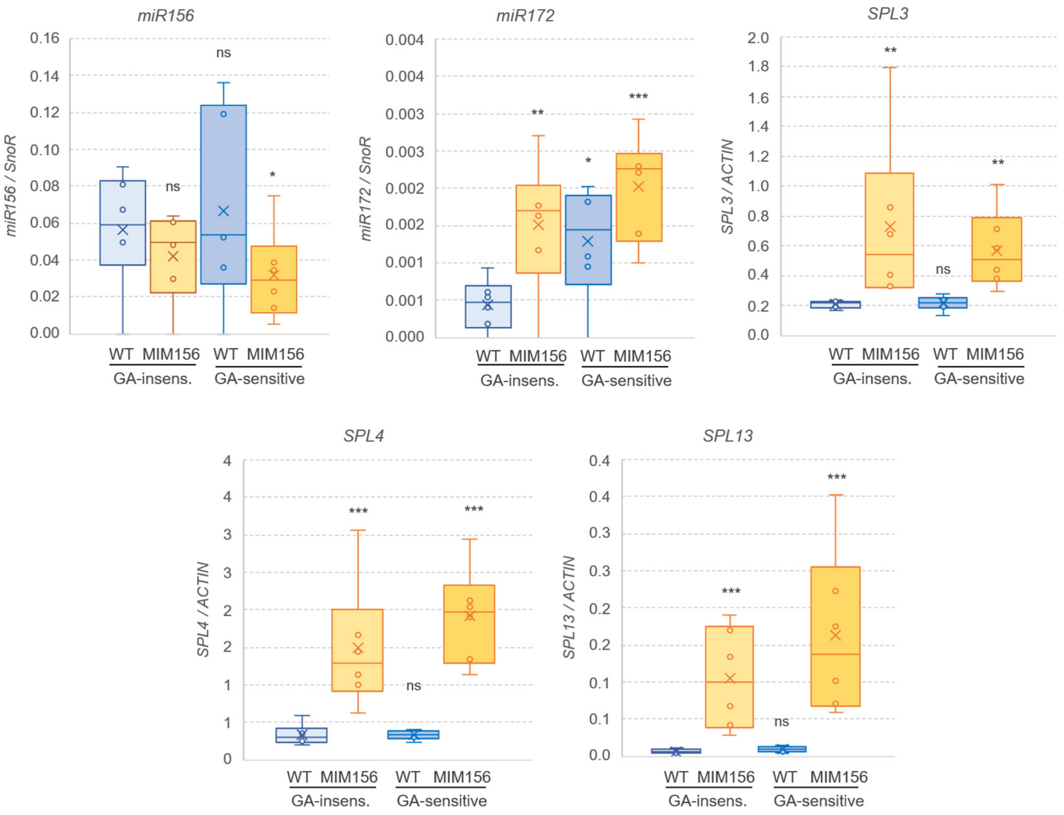
